# Development of multi-species qPCR assays for a stress transcriptional profiling (STP) Chip to assess the resilience of salmonids to changing environments

**DOI:** 10.1101/2024.09.25.615083

**Authors:** Shahinur S. Islam, Daniel D. Heath, Brian Dixon, Phillip Karpowicz, Kelvin Vuu, Jonathon LeBlanc, Nicholas J. Bernier, Kenneth M. Jeffries

**Affiliations:** Great Lakes Institute for Environmental Research, University of Windsor, ON, Canada; Department of Integrative Biology, University of Windsor, ON, Canada; Department of Biology, University of Waterloo, ON, Canada; Department of Biomedical Sciences, University of Windsor, ON, Canada; Department of Integrative Biology, University of Guelph, ON, Canada; Department of Biological Sciences, University of Manitoba, MB, Canada; Department of Animal Science, University of California Davis, CA, USA

**Keywords:** Transcriptomics, Fish health, Climate change, Salmon, Trout, Chars, Whitefishes

## Abstract

Ecologically and socio-economically important salmonid fishes in Canada are threatened by diverse environmental stressors. However, predicting species’ responses to environmental change requires understanding the underlying molecular mechanisms governing environmental stress tolerance. Developing advanced molecular genetic tools will provide opportunities to predict how salmonid fishes will respond to environmental stressors and assess their adaptive potential and vulnerability into the future. Here, we developed a panel of Taqman quantitative PCR (qPCR) assays designed to measure mRNA transcript abundance at selected candidate loci for use across salmonids. We designed and applied those assays for use in a high-throughput nanofluidic OpenArray Stress Transcriptional Profiling Chip (STP-Chip) capable of 2688 simultaneous qPCR at multiple gene loci (112 targets for 12 samples in duplicate). Using the nanofluidic STP-Chip, we tested these 112 multi-species qPCR assays using gill, liver and muscle tissue from eight species of salmonids across four genera. Of the selected 112 assays, 69 assays showed amplification in gill, 64 in liver, and 67 in muscle across all eight salmonid species. The percentage of assays that showed amplification across three tissues varied between genera: In general, *Salmo*, *Oncorhynchus*, and *Salvelinus* species showed a higher success rate than *Coregonus* species. Stress, circadian rhythm, apoptosis, growth-metabolism, and detoxification-relevant assays showed high success rates for amplification across all salmonid species for all three tissues. In contrast, neural plasticity, appetite regulation, osmoregulation, immune function, endocrine disruption, and hypoxia-relevant assays showed low success. Not surprisingly, we observed tissue-specific variation among qPCR amplification patterns. There were significant differences in mRNA transcript abundance among species across the four genera, but we did not see variation between species from the same genus. These qPCR assays can be used to design custom STP-Chips that can be used for quantifying stress in salmonid fish, improving health through more accurate diagnostic tests for disease, and monitoring adaptation to accelerated climate change regionally and globally.

## Introduction

The speed of current global biodiversity loss is unprecedented, and biological responses to the underlying stressors have also been rapid at the species, community, and ecosystem levels (Bernatchez et al. 2023). Accelerated global change is profoundly altering biodiversity at local, regional, and global scales (Heino et al. 2009). At least 200 vertebrate species have become extinct within the last century, and numerous species are facing detrimental consequences to their persistence (Ceballos et al. 2017). Despite huge biodiversity losses across all biomes, an understanding of how taxa will respond to future changing environments remains elusive. Major changes in aquatic environments, such as increased water temperature, decreasing levels of dissolved oxygen, and acidification, among others, will have direct effects on aquatic organisms by challenging their physiological limits (Klein et al. 2022; Sandoval-Castillo et al. 2020), and indirect effects by increasing the spread of novel pathogens (Baker et al. 2022) or by facilitating the introduction of exotic species that may alter competition and predation dynamics in aquatic communities (Gilman et al. 2010). Advanced ‘omics’ technologies provide an opportunity to predict how species will respond to environmental change, identify mechanisms underlying those changes, and assess the adaptive potential and vulnerability of species into the future (Bay et al. 2018; Scheffers et al. 2016).

While most research on the effects of environmental change on biodiversity has concentrated on the terrestrial and marine realms, relatively few studies have investigated these effects on freshwater species (Vörösmarty et al. 2010). Freshwater fish resources in Canada are vast and contribute to Canada’s economy both directly (e.g., recreational and commercial fishing) and indirectly (e.g., ecosystem services) (Howarth et al. 2023). Thriving freshwater fisheries and biodiversity are the lifeblood of many regional and Indigenous communities and have substantial economic, socio-cultural, and ecological value to Canadians (Castañeda et al. 2020). Yet, freshwater fish stocks in Canada are under threat, and to mitigate this, effective management and conservation approaches must be applied (Cooke et al. 2012). Currently, the difficulties of assessing the health and coping capacity of fish in Canada’s > two million lakes and countless rivers are compounded by the limitations of available methods to rapidly screen fish for evidence of physiological stress. Characterizing the health status and potential coping capacity of fishes is therefore crucial for identifying and mitigating increasing freshwater stressors. Thus, there is a need for molecular genetic tools that can rapidly screen across a broad range of organismal responses to multiple stressors in freshwater fishes (Semeniuk et al. 2022).

Salmonid species from the genera *Salmo*, *Oncorhynchus*, *Salvelinus*, and *Coregonus* contribute significantly to the economy through commercial fisheries, recreational fishing, and aquaculture and to the environment through their ecosystem services and biodiversity (Glover et al. 2017). Additionally, salmonids exhibit substantial diversity in their behaviour, morphology, physiology and life history, with patterns of trait variation often replicated within and across species, as well as across different freshwater ecosystems (Bernatchez et al. 2010; Fraser et al. 2011). However, the sustainability of salmonid fish and fisheries in Canada is increasingly affected by climate warming, human-induced habitat alteration, the introduction of aquatic invasive species (AIS) and other drivers of environmental change (Gillis et al. 2024). Salmonid fishes are highly vulnerable to such ecological alterations, resulting in declining populations across freshwater ecosystems in Canada and globally (Reid et al. 2019). Therefore, the conservation and management of salmonid species must evolve to better address their rapidly changing ecosystems.

The development of “omics” (genomics, transcriptomics, metabolomics, etc.) technologies has transformed studies in fish stress physiology over the past decade (López-Maury et al. 2008), by enabling quantification of gene expression processes that mediate an organism’s response to environmental stressors (Schulte 2004). Indeed, gene expression, as estimated by changes in RNA transcript abundance, has been interpreted as putatively representing adaptive variation among natural populations (e.g., Bugg et al. 2023; Wellband et al. 2018). Transcriptional profiling involves targeted quantification of gene transcription at multiple selected gene loci to detect a coordinated multi-locus response to specific stimuli, and this response could be adaptive or maladaptive. As such, transcriptional profiling has applications in assessing health status and adaptive capacity in wild and captive fish populations (Connon et al. 2018). Transcriptional profiling has been proposed for selecting source populations for culture, reintroduction and supplementation for fishery and at-risk species (He et al. 2016). In general, transcriptional profile patterns reflecting phenotypic flexibility can be indicative of an individual’s health status and thus affect population dynamics, species interactions, community ecology, and, ultimately, ecosystem-level processes (Walker et al. 2004).

The goal of this study was to develop 112 multi-species quantitative PCR (qPCR) assays to use with STP-chips to quantify salmonid fishes’ response to environmental stressors (e.g., climate change, AIS, pollution, etc.) that currently affect Canada’s freshwater fisheries and aquaculture. Specifically, we developed a suite of TaqMan® probe-based qPCR assays for 112 genes known to regulate dynamic physiological processes that are key for maintaining homeostasis and health in salmonid fishes. Our study included genes that represent diverse biological functions (e.g., stress response, hypoxia, growth and metabolism, immune function, osmoregulation, apoptosis, detoxification, circadian rhythm, endocrine disruptions, neural plasticity, appetite regulation). These assays will improve fish health assessment and foster better management and conservation of exploited and at-risk salmonid fish stocks not only in Canada but globally. Salmonid STP-Chips will also have important applications in fish culture to screen captive stocks for important performance and production traits (Houston et al. 2020), therefore enhancing production efficiency, sustainability, and profitability.

## Materials and Methods

### Study species and sampling

Eight salmonid species across four genera [*Salmo*: Atlantic salmon (*Salmo salar*), brown trout (*Salmo trutta*); *Oncorhynchus*: Chinook salmon (*Oncorhynchus tshawytscha*), rainbow trout (*Oncorhynchus mykiss*); *Salvelinus*: Brook trout (*Salvelinus fontinalis*), Arctic char (*Salvelinus alpinus*); *Coregonus*: Lake whitefish (*Coregonus clupeaformis*), and bloater (*Coregonus hoyi*)] were selected for this study. Gill, liver and muscle tissue samples from these eight species were collected and stored in high-salt buffer and preserved at -20 ^0^C in the conservation genetics lab at the University of Windsor, Canada. Tissue samples from Atlantic salmon were collected from the Freshwater Restoration Ecology Centre (FREC), University of Windsor. The FREC Atlantic salmon were from the Ontario Ministry of Natural Resources and Forestry (OMNRF) Normandale Fish Culture Station, ON, Canada and were LaHave strain. Tissue samples for brown trout, brook trout and bloater were collected from the Chatsworth Fish Culture Station (OMNRF), ON, Canada. Brown trout were sourced from the Ganaraska River, ON, but were functionally part of the greater Lake Ontario Population. The Brook trout were sourced from Hills Lake Fish Culture Station, ON, Canada and domesticated at Chatsworth Fish Culture Station. Bloater originated from the White Lake Fish Culture Station, ON, Canada. Tissue samples for Chinook salmon were collected from Yellow Island Aquaculture Ltd. BC, Canada. This captive Chinook salmon stock originated from the Department of Fisheries and Oceans Canada (DFO) hatcheries at Robertson Creek and Big Qualicum on Vancouver Island. Tissue samples for rainbow trout, Arctic char and lake whitefish were collected from the Ontario Aquaculture Research Centre (OARC), ON, Canada. The rainbow trout at the OARC were imported from the West Coast of Canada, descendants from brood lines originally maintained by Spring Valley Farm and Blue Springs Farm, BC, Canada. The Arctic char descended from wild gamete collections by DFO, NL, Canada from the Fraser River, Labrador. Arctic char broodstock was developed and distributed by the Rockwood Aquaculture Research Centre in MB, Canada, and arrived at OARC in the mid-1990s. The lake whitefish were descendants of wild-caught fish harvested by commercial fishermen in 2019 near Killarney, Georgian Bay, ON, Canada.

### RNA extraction and cDNA synthesis

Total RNA was extracted from the gill, liver and muscle tissues from a total of 72 fish (n = 8 salmonid species x 3 biological replicates x 3 tissues) using TRIzol (Invitrogen/Life Technologies) with glass mill beads (1mm; BioSpec Products) using a Beadbeater (BioSpec), following the manufacturer’s protocol, and the extracted RNA was stored at -80 ^°^C. RNA integrity was assessed using gel electrophoresis and RNA purity was assessed using A260/280 and A260/230 via NanoQuant for SPARK plate reader (Tecan). Only samples with high purity (A260/280 > 2.0, A260/230 > 1.8) and integrity (clear 18S and 28S RNA bands) samples were used for cDNA synthesis. A total of 2 μg of total RNA was synthesized into c DNA using a High-Capacity cDNA Reverse Transcription (RT) kit (Applied Biosystems, Burlington, ON, Canada). The cDNA was stored at -80 ^°^C until further use.

### Primer and Probe Design and Optimization

One hundred and eight of the 112 candidate genes used in this study (Table 1) were selected based on their role in stress response, hypoxia, immune response, growth and metabolism, detoxification, osmoregulation, neural plasticity, appetite regulation, apoptosis, circadian rhythm, and endocrine disruption. Four endogenous control genes (*rpl7*, *rps9*, *ef1a*, and *rpl13a*) were selected to normalise the transcription profiles of the 108 candidate genes. The primers and probes of the selected genes were designed by the team of the Genomic Network for Fish Identification, Stress and Health project (GEN-FISH; https://gen-fish.ca/). For each candidate gene and endogenous control gene, gene sequences from the four genera of salmonids were downloaded from GenBank (http://www.ncbi.nlm.nih.gov/Genbank/) and aligned using Geneious Prime software v6.1.6. The initial alignments included salmonids (including the sequences available for different subtypes), as well as *Esox lucius* and *Danio rerio* sequences to generate a phylogenetic tree using the Geneious Tree Builder with default settings (i.e., Tamura-Nei Neighbor-joining). The sequence of *Danio rerio* was included as the zebrafish genome is well annotated and helpful for distinguishing subtypes, and the sequence of *Esox lucius* was included as Northern pike is closely related to salmonids and was used as an outgroup to root the phylogenetic tree.

**Table 1:**
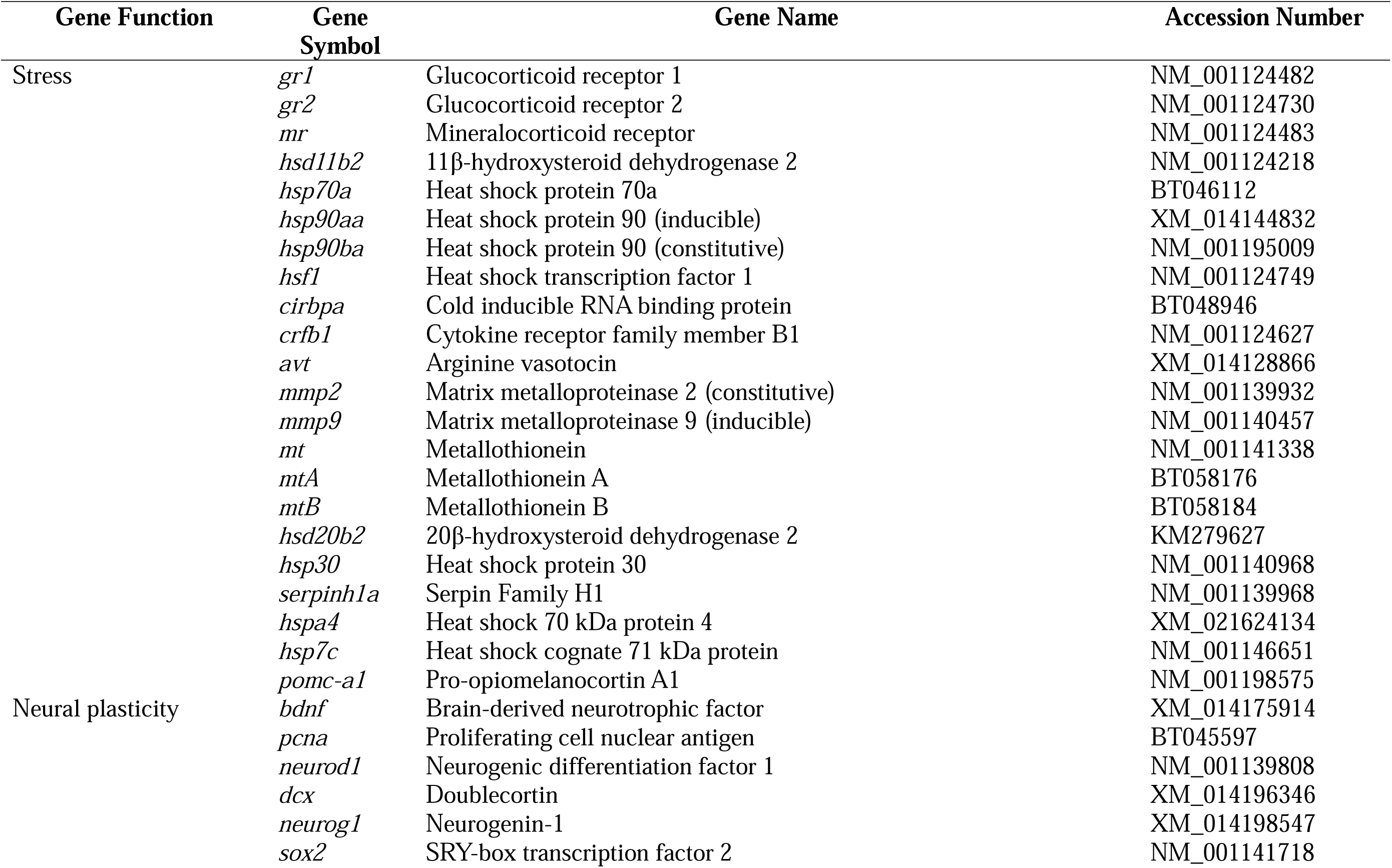
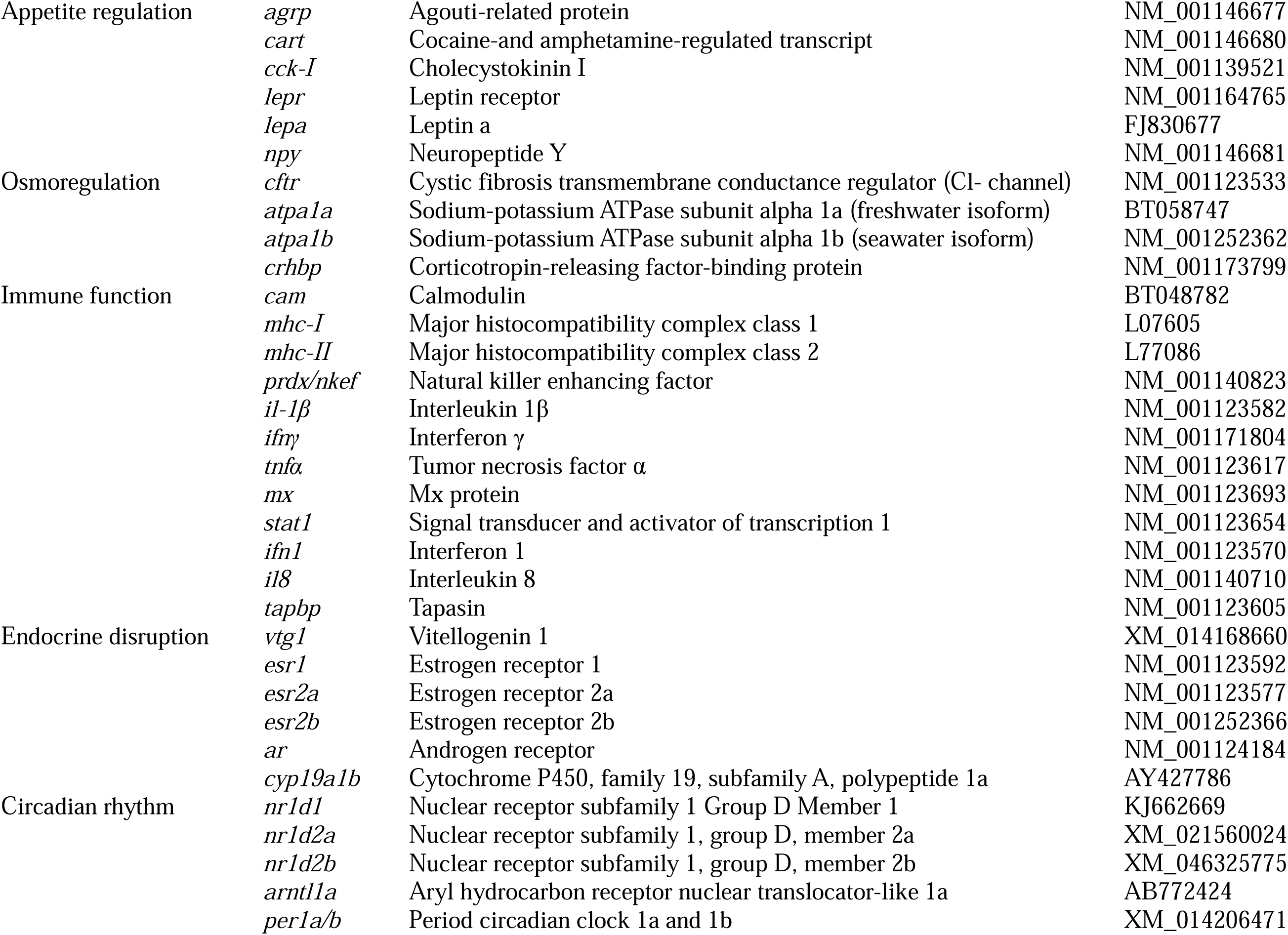
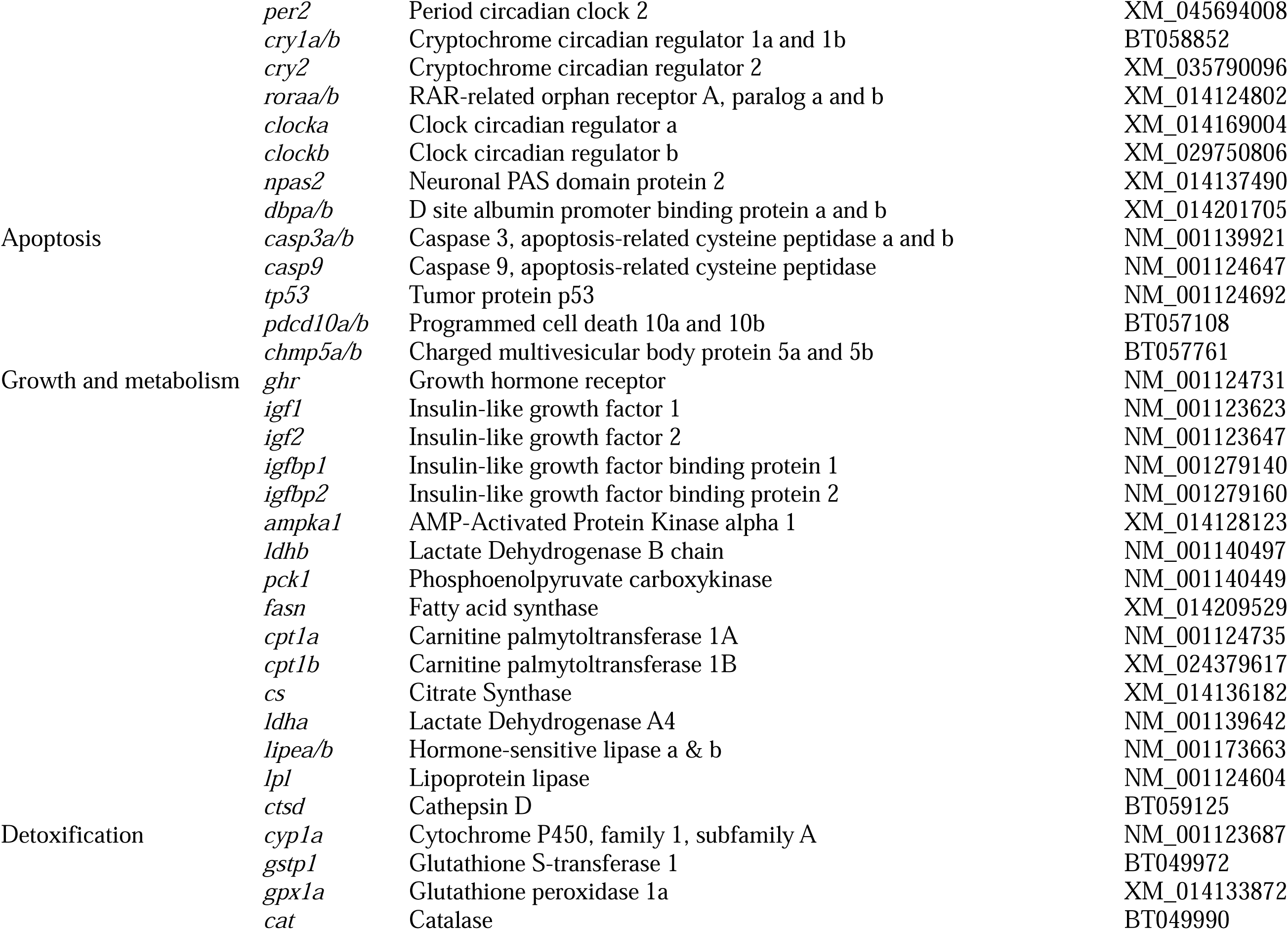
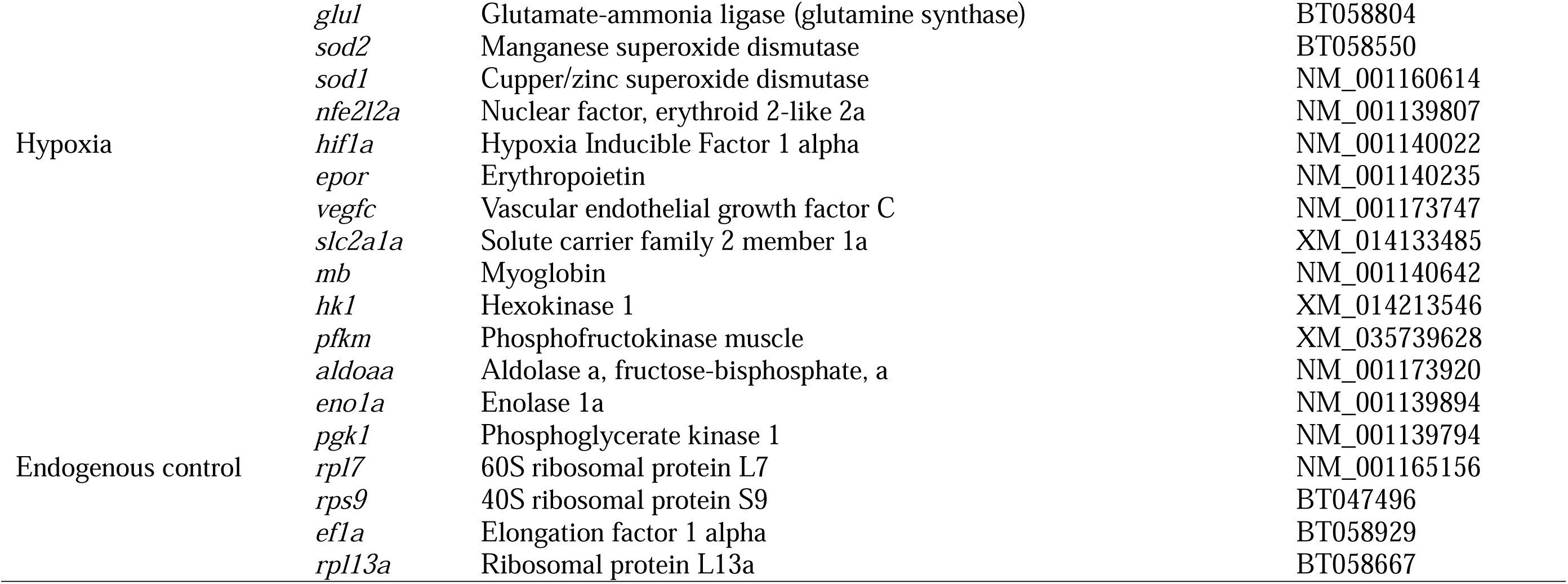
List of 112 selected candidate genes included in the transcriptional profiling panel for salmonids. qPCR assays were developed for all listed genes.

We then highlighted a cluster of sequences that had grouped with the Salmonid orthologs and extracted just the coding region for primer and probe design. The primer and probe sequences were generated based on the consensus sequences. We looked for the primer and probe set that targeted the most conserved portions of the sequence. Finally, we used Primer Express® software (v3.0.1) software to check the parameters (primer/probe Tm, %GC content, primer/probe length, primer/probe composition, amplicon size etc.) for TaqMan® MGB quantification and make the primer/probe compatible with the OpenArray platform (Applied Biosystems, Burlington, ON, Canada).

### Quantitative Real-Time PCR

TaqMan® OpenArray® chips from Applied Biosystems (Burlington, ON, Canada) were used to quantify transcription on a QuantStudio 12K Flex Real Time PCR System following the manufacturer’s protocol. Each chip consists of 24 subarrays of 112 through holes that contained the primer and probe for one of the qPCR assays, resulting in a total of 2,688 qPCR reactions on each chip. Thus 24 cDNA samples were run for each gill, liver and muscle tissue of the 108 target genes along with the 4 endogenous control genes. A 5 µL mixture was prepared for each cDNA sample which contained 2.5 µL TaqMan® OpenArray® Real-Time PCR Master Mix (Applied Biosystems, Burlington, ON, Canada), 1.3 µL ddH_2_O and 1.2 µL cDNA. The 5 µL mixtures were prepared in 384-well plates and loaded into OpenArray® chips using the OpenArray® AccuFill System to reduce the inter-assay variation. A total of 3 OpenArray® chips were used for the 72 cDNA samples. The samples were randomly distributed, and all chips were normalized to the internal reference genes in ExpressionSuite Software v.1.0.3 (Applied Biosystems, Burlington, ON, Canada). We calculated PCR efficiencies for each gene using LinRegPCR (Ramakers et al. 2003). For estimating PCR efficiencies, we exported the raw amplification data from the QuantStudio 12K Flex Real-Time PCR system and separated the imported data by target gene to calculate an individual Window-of-Linearity for each target gene across all samples.

### Statistical analyses

All statistical analyses were performed in R version 4.3.0 (R Core Team 2023). Statistical significance for qPCR data was inferred if *P* < 0.05 after sequential Bonferroni adjustment (Rice 1989). All data were checked visually (Q-Q plot) and statistically (Shapiro-Wilk’s test) for normality, and homoscedasticity was assessed visually (using residuals vs. fitted values) (Crawley 2005). The qPCR datasets for the different tissues (n = 69 for gill tissue, n = 64 for liver tissue, and n = 67 for muscle tissue) were analyzed via principal component analysis (PCA) using the Factoextra R packages (see Figure 3, 4, and 5). Species-specific-driven PCA scores (i.e., PC1 and PC2) and differences in qPCR Δ*C*_T_, were assessed using either ANOVA (parametric) or the Kruskal-Wallis test (non-parametric alternative to one-way ANOVA). Tukey-adjusted multiple comparisons posthoc test were performed for pairwise comparisons between eight salmonid species for each gill, liver and muscle tissue.

## Results

### qPCR confirmation

For gill tissue, sixty-nine assays showed amplification across all eight salmonid species (Figure 1, Supplementary Table 1). Sixty-four assays showed amplification with liver tissue cDNA across all salmonid species (Figure 2, Supplementary Table 2). For muscle tissue, sixty-seven assays showed amplification across the eight salmonid species (Figure 3, Supplementary Table 3). More specifically, for gill tissue, 88% in *S. salar*, 89% in *S. trutta*, 82% in *O. mykiss*, 92% in *O. tshawytscha*, 91% *S. fontinalis*, 86% in *S. alpinus*, and 77% in *C. clupeaformis* and *C. hoyi*; for liver tissue; 83% in *S. salar*, 84% in *S. trutta*, 80% in *O. mykiss*, 86% in *O. tshawytscha*, 91% *S. fontinalis*, 79% in *S. alpinus*, and 73% in *C. clupeaformis* and 71% in *C. hoyi*; and for muscle tissue, 83% in *S. salar*, 84% in *S. trutta*, 80% in *O. mykiss*, 87% in *O. tshawytscha*, 91% *S. fontinalis*, 80% in *S. alpinus*, and 72% in *C. clupeaformis* and 79% in *C. hoyi* showed amplification (Figure 4a, Supplementary Table 4). Assays associated with genes related to stress (gill and liver: 59%, muscle: 68%), circadian rhythm (gill: 69%, liver: 69%, and muscle: 77%), apoptosis (gill, liver, and muscle: 60%), growth and metabolism (gill: 75%, liver: 63%, and muscle: 75%), and detoxification (75% in all tissues) all showed high success rate for amplification and tissue-specific variation across all salmonid species. On the other hand, assays related to neural plasticity (gill: 50%, liver: 33%, and muscle: 17%), appetite regulation (gill: 33%, liver: 17%, and muscle: 17%), osmoregulation (gill: 50%, liver: 25%, and muscle: 25%), immune function (gill: 75%, liver: 58%, and muscle: 42%), endocrine disruption (gill: 17%, liver: 67%, and muscle: 50%), and hypoxia (gill: 50%, liver: 40%, and muscle: 60%) showed overall lower success rates and tissue-specific variation across all salmonid species for all three tissues (Figure 4b, Supplementary Table 5).

**Figure 1.**
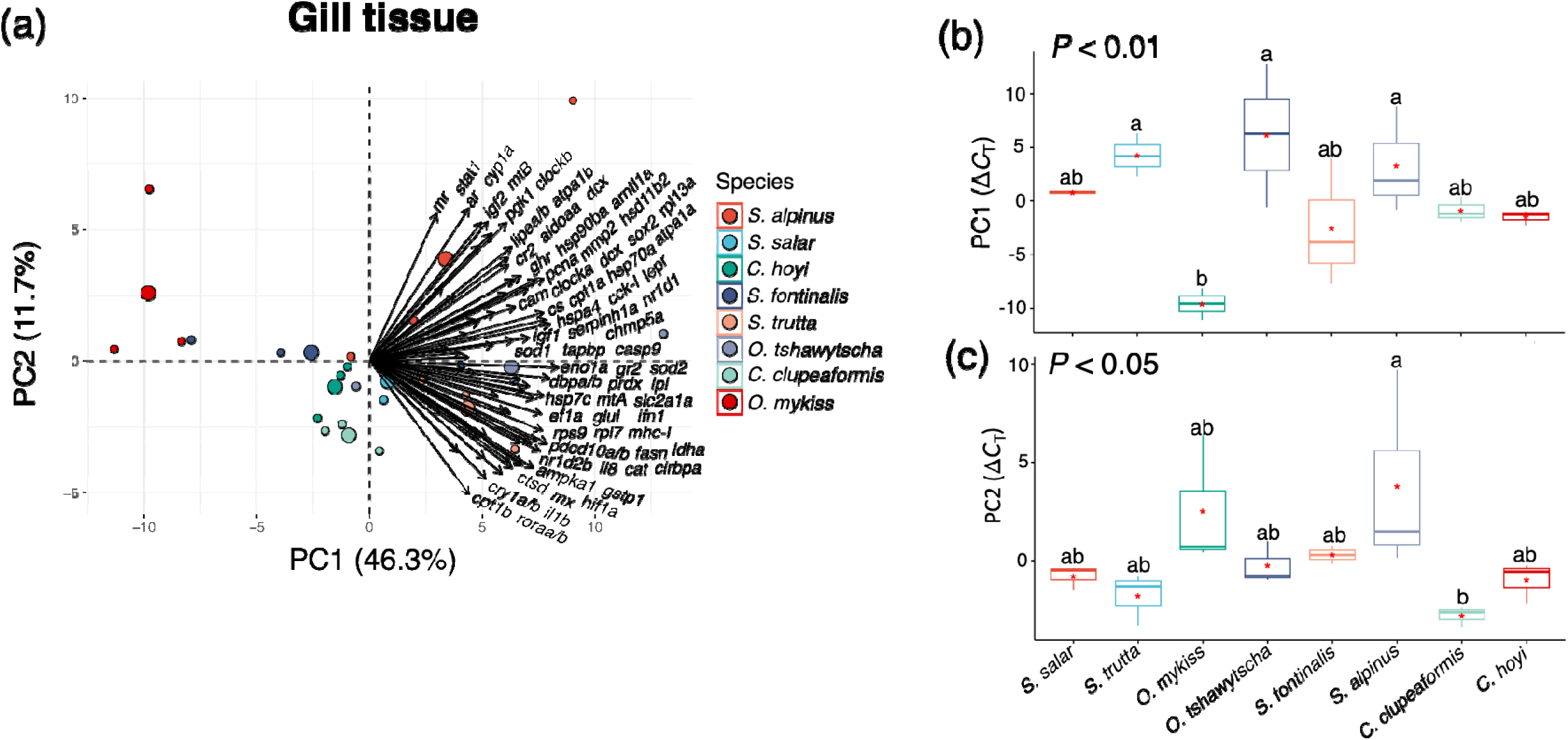
(a-c): Principal component analysis (PCA) of the qPCR assays that showed the mRNA transcript abundance level at gi l tissue. PCA plot illustrating the distribution of all salmonid species included in the study in the multivariate space. Loading vector plot showing the association of each candidate gene of interest with the variance explained by PC1 and PC2. Of the selected 112 assays, forty-three assays at gill tissues did not show amplifications for any of the salmonid species (see Supplementary Table 1); therefore, they were not included in the PCA. The log_2_-transformed Δ*C*_T_ values for all salmonid species were combined to generate PC1 and PC2. Log_2_-transformed Δ*C*_T_ values from all genes for each salmonid species were combined to generate boxplots of PC1 and PC2 scores. Bold line represents median, boxes 25 and 75% quartiles, whiskers 95% confidence interval, dots outliers and red asterisk (*) mean. Different letters denote significant differences in mean Δ*C*_T_ values among eight salmonid species across four genera.

**Figure 2.**
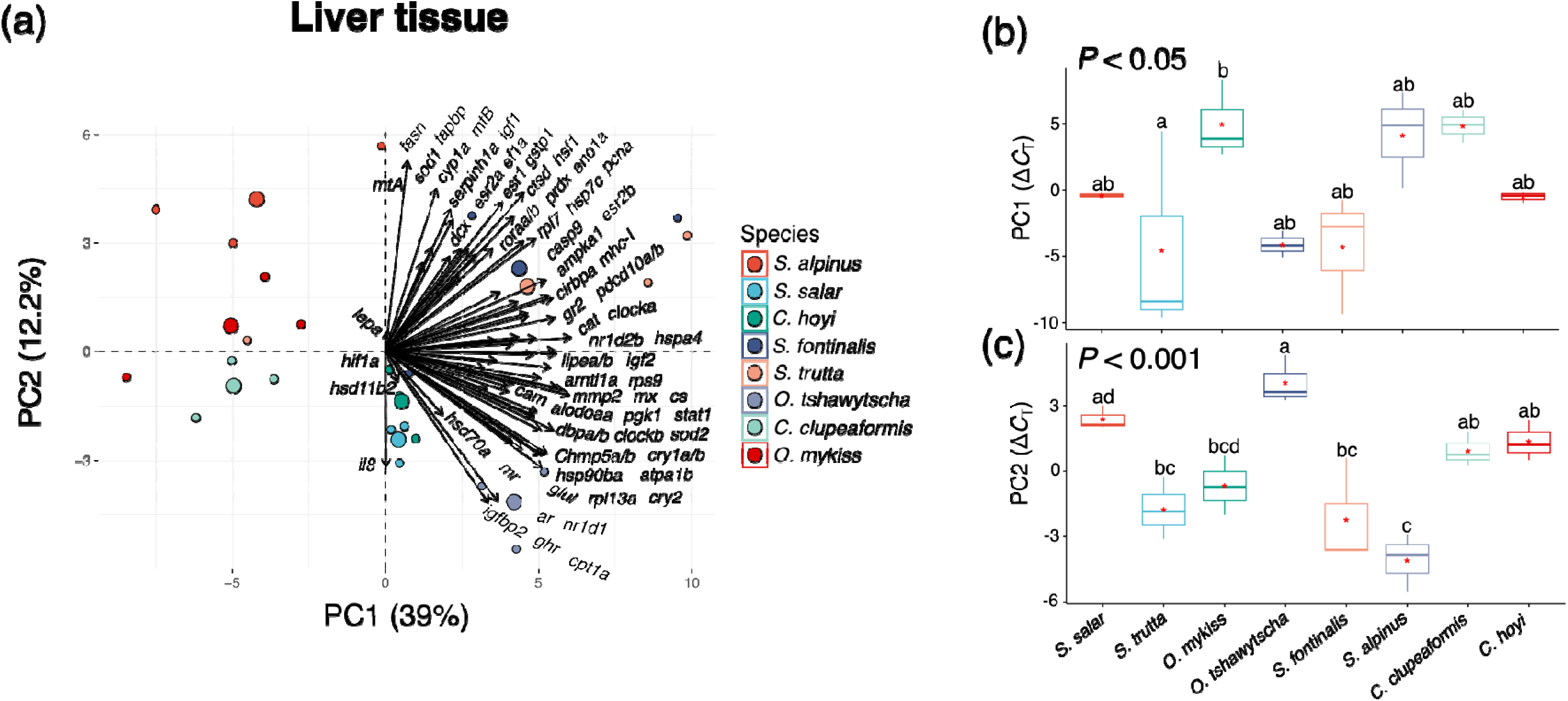
(a-c): Principal component analysis (PCA) of the targeted assays that showed the mRNA transcript abundance level at liver tissue. PCA plot illustrating the distribution of all salmonid species included in the study in the multivariate space. Loading vector plot showing the association of each candidate gene of interest with the variance explained by PC1 and PC2. Of the selected 112 assays, forty-eight assays at liver tissue did not show amplifications for any of the salmonid species (see Supplementary Table 2); therefore, they were not included in the PCA. The log_2_-transformed Δ*C*_T_ values for all salmonid species were combined to generate PC1 and PC2. Log_2_-transformed Δ*C*_T_ values from all genes for each salmonid species were combined to generate boxplots of PC1 and PC2 scores. Bold line represents median, boxes 25 and 75% quartiles, whiskers 95% confidence interval, dots outliers and red asterisk (*) mean. Different letters denote significant differences in mean Δ*C*_T_ values among eight salmonid species across four genera.

**Figure 3.**
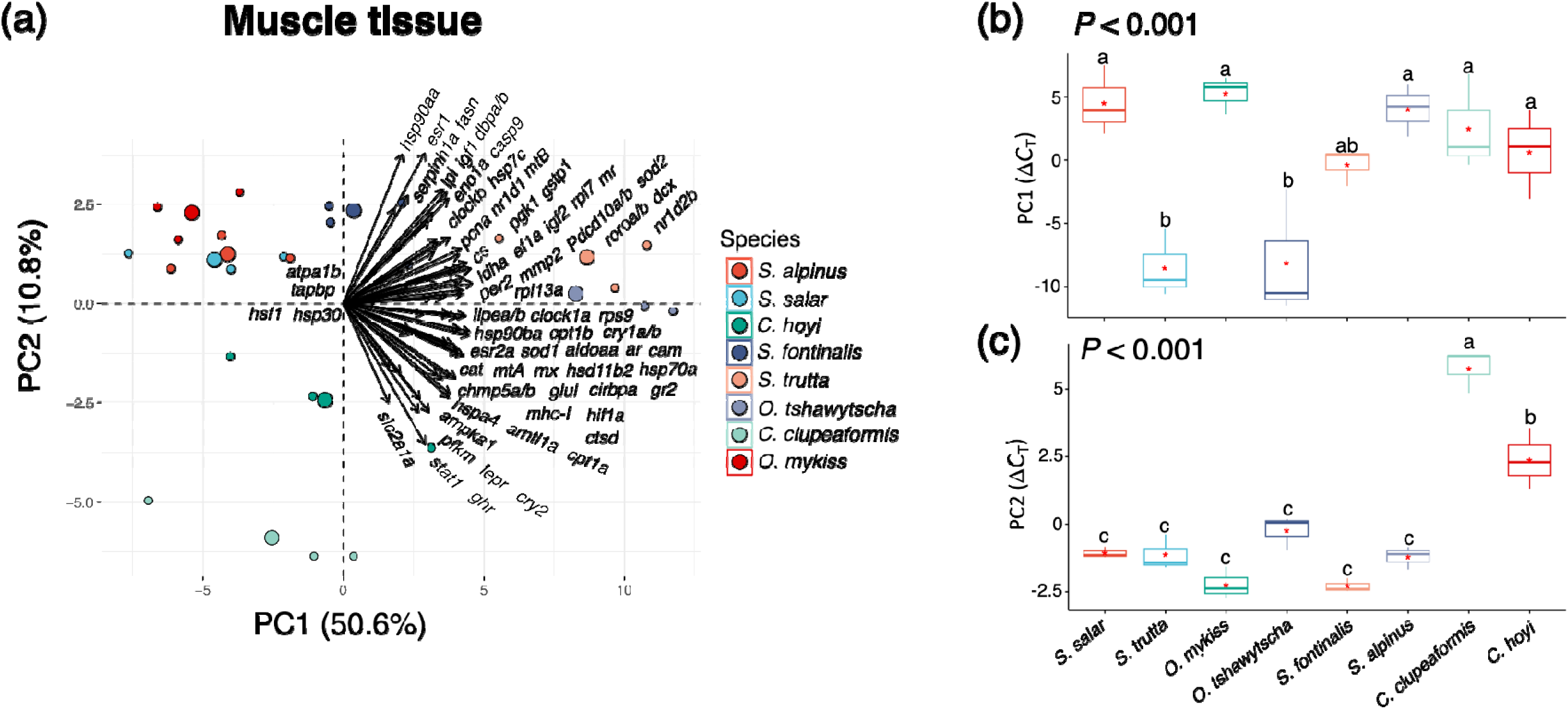
(a-c): Principal component analysis (PCA) of the targeted assays that showed the mRNA transcript abundance level at muscle tissue. PCA plot illustrating the distribution of all salmonid species included in the study in the multivariate space. Loading vector plot showing the association of each candidate gene of interest with the variance explained by PC1 and PC2. Of the selected 112 assays, forty-five assays at muscle tissue did not show amplifications for any of the salmonid species (see Supplementary Table 3); therefore, they were not included in the PCA. The log_2_-transformed Δ*C*_T_ values for all salmonid species were combined to generate PC1 and PC2. Log_2_-transformed Δ*C*_T_ values from all genes for each salmonid species were combined to generate boxplots of PC1 and PC2 scores. Bold line represents median, boxes 25 and 75% quartiles, whiskers 95% confidence interval, dots outliers and red asterisk (*) mean. Different letters denote significant differences in mean Δ*C*_T_ values among eight salmonid species across four genera.

**Figure 4.**
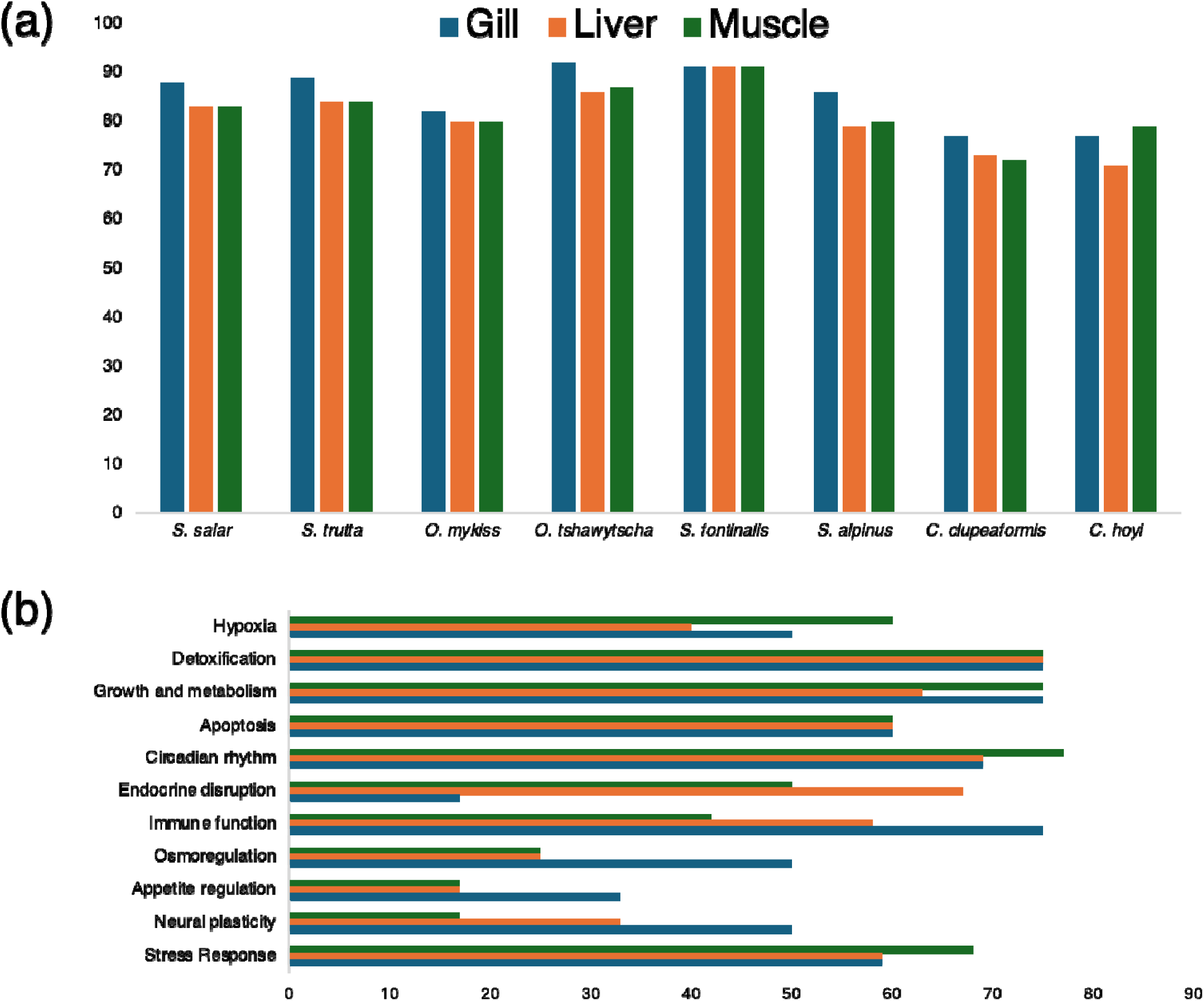
(a-b): (a) Of the 112 assays, the percentage (%) of assays that showed amplification at gill, liver and muscle tissue in all eight salmonid species (for more details, see Supplementary Table 4); (b) the success rate (%) of amplified assays for different biological pathways in gill, liver and muscle tissues (for more details, see Supplementary Table 5).

A PCA for the amplified qPCR genes in gill tissue cDNA (n = 69 assays) revealed species-specific differences along PC1 and PC2 (Figure 1a-c). PC1 and PC2 explained 46.3% and 11.7% of the variance in the mRNA transcript abundance patterns, respectively and showed significant differences among species along PC1 (*P* < 0.01) and PC2 (*P* < 0.05). There was no significant variation in transcript abundance between the two species from same genus along PC1 and PC2 (but see *Oncorhynchus* along PC1). Surprisingly, species from the same genus did not cluster together, with a few exceptions. For example, only species from *Clupeaformis* genus clustered together along with *S. salar* and *S. fontinalis*. Whereas *S. trutta* and *O. tshawytscha* clustered together, but individuals from *O. mykiss* and *S. alpinus* clustered separately. Also, in liver tissue, successful qPCR assays (n = 64) revealed species-specific differences along PC1 and PC2 (Figure 2a-c). PC1 and PC2 explained 39% and 12.2% of the variance in amplified qPCR genes, respectively and showed significant species level differences along PC1 (*P* < 0.05) and PC2 (*P* < 0.001). There was no variation in mRNA transcript levels between the two species from the same genus along PC1 and PC2, with a few exceptions (see *Salmo* and *Oncorhynchus* along PC2). Again, species from the same genus did not cluster together. For example, *S. trutta* and *S. fontinalis*, *S. salar* and *C. hoyi*, and *O. mykiss* and *C. clupeaformis* clustered together, respectively, but individuals from *O. tshawytscha* and *S. alpinus* clustered separately. For muscle tissue cDNA, successfully amplified assays (n = 67) revealed differences across all salmonid species along PC1 and PC2 (Figure 3a-c). PC1 and PC2 explained 50.6% and 10.8% of the variance in functional qPCR assays, respectively and showed significant species-level differences along PC1 (*P* < 0.001) and PC2 (*P* < 0.001). There was little variation in transcript levels between the two species from the same genus along PC1 and PC2 using cDNA from muscle tissue (but see *Salmo* and *Oncorhynchus* along PC1 and *Coregonus* along PC2). For muscle tissue, only species from *Salvelinus* genus clustered together, along with *S. salar* and *O. mykiss*. Species from two different genus formed one cluster (e.g. *S. trutta* and *O. tshawytscha*), but individuals from *C. clupeaformis* and *C. hoyi* clustered separately.

## Discussion

We developed a suite of 112 qPCR assays that can be used to monitor and assess molecular physiological signatures of acute and chronic stress in multiple tissues across diverse salmonid fishes. We found that: (i) of the 112 assays that were developed based on their well-defined biological functions, sixty-nine assays successfully amplified in gill tissue, sixty-four assays in liver tissue, and sixty-seven assays in muscle tissue across eight salmonid species from four different genera; (ii) the percentage of assays that showed amplification across three tissues varied within genus: *Salmo*, *Oncorhynchus*, and *Salvelinus* species showed higher success rate compared to *Coregonus* species; (iii) generally, the stress, circadian rhythm, apoptosis, growth and metabolism, and detoxification related qPCR assays showed higher success rates across the eight salmonid species for all three tissues; whereas, neural plasticity, appetite regulation, osmoregulation, immune function, endocrine disruption, and hypoxia-related qPCR assays showed lower success rate for amplifications across all salmonid species for all three tissues; (iv) there was tissue-specific variation among qPCR success patterns; and (v) there were significant differences in transcript abundance among the four genera but we observed less variation between species within the same genus. These assays will facilitate the design of targeted custom STP-Chips that can be used to provide a quantitative measure of fish health/stress status. Such multifunctional STP-Chips will have applications in commercial fish culture, release and supplementation programs, and for monitoring natural populations.

We aimed to design and test a suite of assays to monitor molecular signatures of acute and chronic stress in fish gills, liver, and muscle tissue. We selected these tissues as they are commonly used for expression profiling in fish, and both gill and muscle biopsies can be used for non-lethal sampling (Jeffries et al. 2021). Gill tissue was chosen for this study as the gill is a multifunctional organ (e.g., with a role in respiration, osmoregulation, acid-base balance, and ammonia excretion). The gill also plays an important role as an immune barrier and has a limited role in detoxification activity, which makes the gill an excellent target for monitoring the transcriptional response of fish to various environmental stressors and pathogens (Akbarzadeh et al. 2018). Liver tissue was chosen as it is a key tissue involved in the processing and storage of nutrients, the synthesis of key enzymes and hormones involved in regulating growth and metabolism, and the metabolism of toxicants (Nikopoulou et al. 2023). Fish muscle is also a heterogeneous tissue that contains different cell types, including myocytes, adipocytes, and fibroblasts (Johnston et al., 2011). Thus, muscle is an important tissue for monitoring fish growth and metabolism, impacts on swim performance and exercise, and apoptosis pathways. This study observed tissue-specific variation in mRNA transcript abundance among qPCR amplification patterns. Specifically, osmoregulation and immune function-relevant assays showed a high success rate in terms of amplification in gill tissue compared to liver and muscle tissues, but it was quite the opposite for endocrine disruption pathways. However, variation in tissue-specific gene transcript response among natural fish populations is well documented (Allendorf et al. 1983; Whitehead and Crawford 2005).

In this study, 62% of assays showed amplification in gill tissue, 57% showed amplification in liver tissue, and 60% showed gene expression in muscle tissue across all eight salmonid species. There was variation in amplification based on different biological functions/pathways. Several important pathways such as stress response, circadian rhythm, apoptosis, growth and metabolism, and detoxification-relevant assays showed high success rates for amplification across all eight salmonid species for gill, liver, and muscle tissue, but some osmoregulatory function pathways showed lower success rates for amplifications across all tissues. This may reflect the endogenous level of expression of these genes, with ones that have higher constitutive expression across tissues being readily detected (i.e., growth and metabolism), while others that are expressed by a subpopulation of tissue cells or context-dependent (i.e., endocrine disruption) harder to detect. In this study, we aimed to design multi-species qPCR assays for producing custom STP-Chips designed to include tissue-specific expressed genes (e.g., thermal stress/hypoxia chip for gill, heart, and muscle, osmoregulation chip for gill, immune chip for liver, head kidney, and spleen, and neural function and appetite regulation chip for brain, etc.). It is also important to note that all fish used in this study did not have a stressor exposure applied before sampling, i.e., we are reporting transcriptional profiles of resting fish in ambient conditions (although they could still have been stressed before sampling). As such, we have no *a priori* reason to expect hypoxia-responsive genes, cellular stress-responsive genes, immune genes, detoxification genes, apoptosis-related genes, etc., to have been expressed at high levels in these fish. Therefore, our results are conservative because expression of the majority of the genes selected for the STP-chips would be upregulated in stressed fish. Thus, studies of challenged fish will likely show a much higher percentage of detectable transcription.

In this study, some assays did not show amplification in *Coregonus* species while they did in *Salmo*, *Oncorhynchus*, and *Salvelinus* species. During each assay design, sequences that showed higher identities/similarities from all four salmonid genera were included in multiple sequence alignments. Although we designed assays at the most conserved portions of the sequences, multiple alignments of all species showed that *Coregonus* sequences were more divergent than the other salmonid genus, which might reflect on the lack of amplification for *Coregonus* species (Lecaudey et al. 2018). Also, we detected significant association in transcript abundance among eight species across four genera but did not find differences between species from the same genus (with a few exceptions), which is expected as salmonids exhibit remarkably high levels of genetic and trait variation across species (Bernatchez et al. 2010).

Salmonid fishes play a vital economic, ecological, and societal role in Canada and internationally. Yet, due to anthropogenic influences such as climate change, introduction of AIS, and habitat destruction, salmonid fish species are under threat globally. To mitigate these threats, salmonid populations need effective management and conservation approaches. Currently, the logistic difficulties of monitoring salmonid population in Canada are compounded by the limitations of conventional (e.g., capture-based) methods. Under the umbrella of GEN-FISH, the transcriptomic approaches we developed would help address questions regarding salmonid population and overall ecosystem health. Further, tools to rapidly screen wild fish for signs of the effects of environmental stress can provide insight into whether the populations can cope with environmental stressors such as climate change and invasive species. Future development of salmonid STP-chips will aid in assessing salmonid fish health and environmental coping capacity. Managers and conservationists will be able to use custom STP-chips to measure stress and coping responses of salmonids to single- and multiple-stressor combinations across a diversity of biological scales. Multi-scale laboratory and field validation of those custom chips will generate useful tools for real-world applications that both monitor exposure to key environmental stressors, and estimate the impacts of these responses on salmonid performance and fitness.

## Acknowledgements

The authors would like to thank P. Voyer, K. Patel, Dr. J. Jeffrey, T. Mackey, S. Mackie, and Dr. Audet lab for technical assistance. The GEN-FISH project is funded by a Genome Canada Large Scale Applied Research Project.

## Author Contributions

SSI contributed to assay design, completed the lab work, data analyses and manuscript preparation. KMJ and DDH contributed to study design, data analyses and manuscript preparation. NJB, BD, PK, KV and JL contributed to manuscript preparation and revision.

## Data Availability

Data from qRT PCR assays will be made available upon publication.

