## Supplementary Files for "Development of multi-species qPCR assays for a stress transcriptional profiling (STP) Chip to assess the resilience of salmonids to changing environments"

Supplementary Table 1: TaqMan primer and probe sequences, PCR efficiencies, and mean C_T_ levels for the 112 candidate genes for gill tissue across eight salmonid species

| Biological function | Gene symbol | Primer and probe sequences | Efficiency (%) | C_T_ level | | | | | | | |
| --- | --- | --- | --- | --- | --- | --- | --- | --- | --- | --- | --- |
|  |  |  |  | *S. salar* | *S. trutta* | *O. mykiss* | *O. tshawytscha* | *S. fontinalis* | *S. alpinus* | *C. clupeaformis* | *C. hoyi* |
| Stress | *gr1* | F: GTCTTTGGCCTGTATCCCCC | 107 | 26.17 | 25.52 | 26.87 | 25.36 | - | - | 26.14 | 27.13 |
|  |  | R: AGCTCGACATCCCTGATCCA |  |  |  |  |  |  |  |  |  |
|  |  | P: TGCCCTCGGTCAGTGA |  |  |  |  |  |  |  |  |  |
|  | *gr2* | F: ATGGCAGACCAGTGTGAACAGAT | 124.5 | 26.84 | 25.84 | 27.55 | 25.06 | 26.55 | 25.58 | 25.31 | 25.92 |
|  |  | R: GAGCAGCAGCAGAACCTTCAT |  |  |  |  |  |  |  |  |  |
|  |  | P: AGACTGCAGGTGTCTCA |  |  |  |  |  |  |  |  |  |
|  | *mr* | F: CATAGTCAATGTCAGCTGCTCCC | 111.7 | 26.49 | 25.26 | 24.81 | 25.14 | 26.78 | 24.56 | 25.50 | 25.99 |
|  |  | R: GCTGCTGCTCTGGCTTCTTCT |  |  |  |  |  |  |  |  |  |
|  |  | P: CCAGCAGCAACACA |  |  |  |  |  |  |  |  |  |
|  | *hsd11b2* | F: CTGTCTAGCAGCGTACGGAGC | 105.6 | 26.03 | 25.87 | 26.32 | 25.45 | 27.28 | 25.17 | 26.53 | 27.50 |
|  |  | R: ATGGTGGACACTTTGACCCC |  |  |  |  |  |  |  |  |  |
|  |  | P: CCTGTTCATCAACACACT |  |  |  |  |  |  |  |  |  |
|  | *hsp70a* | F: GAGAACACTGTCCTCCAGCTCC | 112.7 | 25.79 | 27.68 | 28.60 | 25.63 | 26.90 | 27.31 | 28.38 | 29.12 |
|  |  | R: CCCTGAAGAGGTCGGAACAC |  |  |  |  |  |  |  |  |  |
|  |  | P: ACACCTCCATCACCAGG |  |  |  |  |  |  |  |  |  |
|  | *hsp90aa* | F: AAGATCGAGGTCACCCCTGA | 96.7 | 29.83 | 30.68 | - | - | 29.87 | - | - | 34.04 |
|  |  | R: GGTGCCAGACTTTGCAATGG |  |  |  |  |  |  |  |  |  |
|  |  | P: CGGCATCGGCATGA |  |  |  |  |  |  |  |  |  |
|  | *hsp90ba* | F: GAGGTGGAGGAGGACGAGTACA | 85.6 | 26.08 | 26.16 | 29.42 | 26.54 | 26.43 | 24.15 | 26.79 | 27.10 |
|  |  | R: GCTGTGAAGTGGATGTGGGA |  |  |  |  |  |  |  |  |  |
|  |  | P: TCTCCAGGGACACAGAC |  |  |  |  |  |  |  |  |  |
|  | *hsf1* | F: CCCAAGTTCAGCAGGCAGTAC | 129.7 | 25.27 | 24.20 | 25.74 | 24.98 | 25.36 | 24.64 | 24.08 | 26.37 |
|  |  | R: GCCGTGAAGAGACCGGTACT |  |  |  |  |  |  |  |  |  |
|  |  | P: TGCAGGGCTCGCCT |  |  |  |  |  |  |  |  |  |
|  | *cirbpa* | F: CGGGAAGGTCTCGTGGATT | 128 | 21.60 | 20.87 | 24.28 | 21.45 | 22.29 | 21.36 | 20.40 | 21.29 |
|  |  | R: TGGTTCTGCCATCGACAGACT |  |  |  |  |  |  |  |  |  |
|  |  | P: CATTGGAGGGAATGAA |  |  |  |  |  |  |  |  |  |
|  | *crfb1* | F: TCATTGCTTTCTTACCGCGC | - | - | - | - | - | - | - | - | - |
|  |  | R: AGGAGGGGAAGAGACTGTTGCT |  |  |  |  |  |  |  |  |  |
|  |  | P: CCGCTCCACATCACGA |  |  |  |  |  |  |  |  |  |
|  | *avt* | F: CTAGACCCAGACTGCCTAGAGGAC | 99.1 | 30.56 | 31.19 | - | 29.62 | 30.17 | 30.79 | - | - |
|  |  | R: GCCAAACCACCCATTAAGGC |  |  |  |  |  |  |  |  |  |
|  |  | P: ACGTCAGTCACCCAGCGA |  |  |  |  |  |  |  |  |  |
|  | *mmp2* | F: CGCTGTGGAGTTCCTGATGTT | 107 | 23.91 | 22.65 | 25.05 | 22.21 | 24.10 | 22.66 | 25.21 | 23.95 |
|  |  | R: AGGTCAGGAGAGTGGCCTAGAA |  |  |  |  |  |  |  |  |  |
|  |  | P: AGGAAACCCAAGTGGCA |  |  |  |  |  |  |  |  |  |
|  | *mmp9* | F: ACGGCAAAGCATGTGTGTTC | 129.3 | 24.75 | 26.44 | 26.73 | - | 26.49 | 22.96 | - | - |
|  |  | R: GGAACACCAGCGGTATCCAT |  |  |  |  |  |  |  |  |  |
|  |  | P: AGGTTGCACGACGGAA |  |  |  |  |  |  |  |  |  |
|  | *mt* | F: GCTCAAAAACTGGACGCTGC | 118.1 | 27.57 | - | - | 29.14 | - | - | - | - |
|  |  | R: GGCAGCAGGAACAACAACTTT |  |  |  |  |  |  |  |  |  |
|  |  | P: TACAAACTGCGGATGTGC |  |  |  |  |  |  |  |  |  |
|  | *mtA* | F: TGGATCCTTGTGAATGCTCCA | 104.7 | 24.78 | 23.80 | 25.55 | 24.57 | 24.72 | 24.74 | 24.72 | 25.36 |
|  |  | R: CTTACAACTGGTGCATGCGC |  |  |  |  |  |  |  |  |  |
|  |  | P: CGGTGGATCCTGCAAG |  |  |  |  |  |  |  |  |  |
|  | *mtB* | F: GAAAAGTTGCTGCCCCTGC | 111.2 | 25.57 | 24.02 | 25.36 | 23.84 | 25.49 | 22.97 | 26.97 | 26.20 |
|  |  | R: AACAGCTGGTATCGCAGGTCTT |  |  |  |  |  |  |  |  |  |
|  |  | P: TTCAGGCTGTGTGTGCA |  |  |  |  |  |  |  |  |  |
|  | *hsd20b2* | F: ACATTGTACTGGTCAGCCGGT | 112.3 | 26.62 | 24.28 | - | 26.54 | 24.41 | 23.86 | 27.18 | 25.31 |
|  |  | R: TGGCCCTCTGTGAAGTCTGTC |  |  |  |  |  |  |  |  |  |
|  |  | P: CCAACATGGACGCCAGA |  |  |  |  |  |  |  |  |  |
|  | *hsp30* | F: TGCTGTGTTCCCGAGGATTC | 87.1 | 29.01 | 26.58 | - | - | - | - | 33.73 | - |
|  |  | R: TCTCTGCAGTAGATCCCGCTG |  |  |  |  |  |  |  |  |  |
|  |  | P: TGTGCGCAGTCTAT |  |  |  |  |  |  |  |  |  |
|  | *serpinh1a* | F: AGCGCTGTGAAGTCCATCAA | 104.4 | 24.06 | 22.86 | 26.45 | 24.38 | 24.03 | 23.30 | 26.30 | 26.02 |
|  |  | R: TGATGATCATGGCCCCATC |  |  |  |  |  |  |  |  |  |
|  |  | P: CGGCCAAGTCCACCGA |  |  |  |  |  |  |  |  |  |
|  | *hspa4* | F: ACTGCTGAGACCGCAATGAA | 120.5 | 24.32 | 24.82 | 27.17 | 24.11 | 25.43 | 24.28 | 25.28 | 25.51 |
|  |  | R: TGCGTCAGTGTAGAAGCTGGG |  |  |  |  |  |  |  |  |  |
|  |  | P: ACCTGTGGCTGATTGT |  |  |  |  |  |  |  |  |  |
|  | *hsp7c* | F: TAGCCAACGACCAGGGAAAC | 111.3 | 22.63 | 23.35 | 27.41 | 23.09 | 25.46 | 25.95 | 27.63 | 27.86 |
|  |  | R: CGTCCCCAATCAGCCTTTCT |  |  |  |  |  |  |  |  |  |
|  |  | P: ACCCAGTTATGTCGCCTT |  |  |  |  |  |  |  |  |  |
|  | *pomc-a1* | F: AGAACAGCATCCTGGAGTGCA | - | - | - | - | - | - | - | - | - |
|  |  | R: GGGAGTTGGGTTGGAGATGG |  |  |  |  |  |  |  |  |  |
|  |  | P: ACCTCACCGCCGAAT |  |  |  |  |  |  |  |  |  |
| Neural plasticity | *bdnf* | F: TCCTTCCTCTGTGGTCACCAC | 110 | 30.14 | 30.36 | - | 30.44 | 31.35 | 29.26 | 30.96 | 31.68 |
|  |  | R: AGGCACTTGGTTGCTGATCA |  |  |  |  |  |  |  |  |  |
|  |  | P: TGTCGACCTGTATGCAT |  |  |  |  |  |  |  |  |  |
|  | *pcna* | F: TGGAGGCTCTGAAGGACCTG | 110.3 | 24.02 | 24.11 | 25.85 | 23.49 | 24.68 | 23.48 | 24.85 | 25.09 |
|  |  | R: GAGCTGAACTAGGGAGACGTGG |  |  |  |  |  |  |  |  |  |
|  |  | P: CTGTTGGGACGTGAGCT |  |  |  |  |  |  |  |  |  |
|  | *neurod1* | F: ATGTCATCAGACCCCCAGGA | 111.9 | 36.20 | 30.78 | - | 31.37 | 31.33 | - | - | - |
|  |  | R: GATTCGAGTGCGCCGTTTT |  |  |  |  |  |  |  |  |  |
|  |  | P: CGACCTGGACTGATGA |  |  |  |  |  |  |  |  |  |
|  | *dcx* | F: GTCTGTGAACGTGAAGGCCTC | 102.2 | 29.84 | 27.96 | 29.96 | 29.17 | 27.74 | 27.90 | 29.75 | 28.76 |
|  |  | R: TTGGGCCGAACAAACTCCT |  |  |  |  |  |  |  |  |  |
|  |  | P: AGCCAGAAGAACCTG |  |  |  |  |  |  |  |  |  |
|  | *neurog1* | F: AGAGCGAGGAATGAAGCCAC | 83.9 | - | - | - | 29.91 | - | - | 31.49 | - |
|  |  | R: GTTTCTCTCGCGGTCGTTTG |  |  |  |  |  |  |  |  |  |
|  |  | P: CGTGGTCAAGAAGAA |  |  |  |  |  |  |  |  |  |
|  | *sox2* | F: TCAGACAGGAGACCTACGGGAC | 114.6 | 27.31 | 24.89 | 27.89 | 25.22 | 25.57 | 27.04 | 26.93 | 26.56 |
|  |  | R: TTGGGACATGTGGAGTCTGC |  |  |  |  |  |  |  |  |  |
|  |  | P: TTACCGGGCGCTGAG |  |  |  |  |  |  |  |  |  |
| Appetite regulation | *agrp* | F: ATACTGCTGGACCCTGTGCCT | - | - | - | - | - | - | - | - | - |
|  |  | R: CGGGATGGGAGTCGTCTAGA |  |  |  |  |  |  |  |  |  |
|  |  | P: CCTGATCCAGCTGGCTA |  |  |  |  |  |  |  |  |  |
|  | *cart* | F: TGACCAACGAAAAGCAACTGC | 101.7 | - | - | - | 29.68 | 29.34 | - | - | - |
|  |  | R: AGGGACTTGGCCGAATTTCT |  |  |  |  |  |  |  |  |  |
|  |  | P: CTGCAGACCAAGAGA |  |  |  |  |  |  |  |  |  |
|  | *cck-I* | F: GGGTCCCAGCCACAAGATAA | 104.5 | 29.04 | 30.01 | 31.02 | 29.97 | 30.65 | 29.71 | 31.02 | 29.94 |
|  |  | R: CTCGTACTCCTCTGCACTGCG |  |  |  |  |  |  |  |  |  |
|  |  | P: TGGGCTGGATGGAC |  |  |  |  |  |  |  |  |  |
|  | *lepr* | F: CGCTGTATGCACATCAACGG | 101.7 | 30.02 | 29.09 | 29.38 | 28.15 | 30.54 | 27.87 | 27.41 | 29.22 |
|  |  | R: GTCAGGAGCTCTGCTGTTGTGA |  |  |  |  |  |  |  |  |  |
|  |  | P: CCGGGACCTGGAGTGA |  |  |  |  |  |  |  |  |  |
|  | *lepa* | F: AAGCCTGCCTTCCATAGTGGA | 121.9 | 31.77 | - | 26.66 | - | 28.14 | 30.72 | 29.73 | - |
|  |  | R: TGAGGTCTGCCCAGTCTAGGA |  |  |  |  |  |  |  |  |  |
|  |  | P: TGGGATTCTACCAGGACC |  |  |  |  |  |  |  |  |  |
|  | *npy* | F: AACCTCATCACAAGGCAGAGGT | 85.3 | 30.00 | - | - | 29.95 | 28.14 | - | - | - |
|  |  | R: ACGTGTCTGTGCTCTCCTTCAG |  |  |  |  |  |  |  |  |  |
|  |  | P: AGGTCCAGCCCTGAC |  |  |  |  |  |  |  |  |  |
| Osmoregulation | *cftr* | F: CAGGCAGAAAAGAGCAGGGA | 126 | 25.55 | 24.90 | 27.89 | 23.65 | 26.41 | 27.15 | - | - |
|  |  | R: TCTCCATCACCTCCTCCCAG |  |  |  |  |  |  |  |  |  |
|  |  | P: TCGTCGCCTGGCTC |  |  |  |  |  |  |  |  |  |
|  | *atpa1a* | F: GGTGGTTGGAGATCTGGTGG | 103.8 | 21.34 | 21.48 | 22.90 | 20.09 | 22.10 | 20.81 | 19.39 | 21.58 |
|  |  | R: CACCAGTGAGGGAGGAGTTGTC |  |  |  |  |  |  |  |  |  |
|  |  | P: ATTTGCGTATTGTCTCTG |  |  |  |  |  |  |  |  |  |
|  | *atpa1b* | F: GGATGCTTGGCTGGGATTT | 102.2 | 21.41 | 21.88 | 22.59 | 21.36 | 21.57 | 21.16 | 21.91 | 21.16 |
|  |  | R: CTCTGTCCTGGTAGGTGGGTTT |  |  |  |  |  |  |  |  |  |
|  |  | P: CAGGCCCTGCTGCT |  |  |  |  |  |  |  |  |  |
|  | *crhbp* | F: GAGCCCAACCAGGTCATCAA | 103.2 | 32.01 | 28.20 | 31.04 | 30.36 | 29.05 | - | 30.32 | 30.09 |
|  |  | R: CTCTCCCTTCATCACCCAGC |  |  |  |  |  |  |  |  |  |
|  |  | P: CGTTGACATCGACTGCA |  |  |  |  |  |  |  |  |  |
| Immune function | *cam* | F: GAGGAGCAGATTGCCGAGTT | 122.8 | 22.80 | 22.67 | 24.63 | 21.57 | 22.67 | 21.10 | 22.84 | 21.97 |
|  |  | R: CATGACAGTGCCCAGCTCTTT |  |  |  |  |  |  |  |  |  |
|  |  | P: CGCTCTTTGACAAGGA |  |  |  |  |  |  |  |  |  |
|  | *mhc-I* | F: AGTCCCTCCCTCAGTGTCTCTG | 111.2 | 19.63 | 18.38 | 21.75 | 17.38 | 20.98 | 19.49 | 18.66 | 18.90 |
|  |  | R: AGGACACCATGACTCCACTGG |  |  |  |  |  |  |  |  |  |
|  |  | P: AGTGACCTGCCACGCG |  |  |  |  |  |  |  |  |  |
|  | *mhc-II* | F: CTCACAGCAGCATCTACCCCA | 103.2 | 28.81 | 28.51 | 24.51 | 21.66 | 26.24 | - | 21.88 | 22.99 |
|  |  | R: CACCTGACTCTGACAGGTGCAG |  |  |  |  |  |  |  |  |  |
|  |  | P: TGGAGAACACCCTCATCT |  |  |  |  |  |  |  |  |  |
|  | *prdx* | F: GGATCAACACCCCCAGGAA | 112.3 | 23.40 | 22.46 | 25.67 | 22.72 | 23.77 | 22.74 | 23.93 | 24.87 |
|  |  | R: GTCCTCCTTCAGCACTCCGTA |  |  |  |  |  |  |  |  |  |
|  |  | P: CCCTTGTGGCTGACCT |  |  |  |  |  |  |  |  |  |
|  | *il-1β* | F: TCAGGGTCTGGATCTGGAGG | 88.3 | 25.99 | 25.89 | 28.62 | 26.04 | 27.89 | 28.52 | 27.68 | 27.67 |
|  |  | R: CTCGGTTCCCATGGTAACCC |  |  |  |  |  |  |  |  |  |
|  |  | P: ACCCCATCACCATGC |  |  |  |  |  |  |  |  |  |
|  | *ifnγ* | F: TGTTTTCCCCAAGGACACGT | 126.7 | 30.16 | 28.39 | 28.87 | 25.54 | - | 28.63 | 31.78 | - |
|  |  | R: CCGATACACGTCCAGAACCA |  |  |  |  |  |  |  |  |  |
|  |  | P: TGAGCGGAGGGTGTT |  |  |  |  |  |  |  |  |  |
|  | *tnfα* | F: CTCAACTCTGTACGCACCGTG | 113.5 | 28.45 | 29.48 | 29.58 | 27.89 | 28.31 | 29.24 | - | - |
|  |  | R: TCTCCCTTCTCCAGGCTGAA |  |  |  |  |  |  |  |  |  |
|  |  | P: CGAGGCTGCGAGTGA |  |  |  |  |  |  |  |  |  |
|  | *mx* | F: GGAGGAGATTGAGGACCCCT | 98.4 | 25.88 | 25.33 | 27.44 | 23.37 | 27.43 | 26.34 | 24.46 | 25.85 |
|  |  | R: ATCACTGATACCCACCCCCA |  |  |  |  |  |  |  |  |  |
|  |  | P: TGAAGCCCAGGATGAA |  |  |  |  |  |  |  |  |  |
|  | *stat1* | F: CAGAAAGGCTTCCTGGAGGG | 105.5 | 24.55 | 23.61 | 24.77 | 24.30 | 24.19 | 22.94 | 23.90 | 23.90 |
|  |  | R: CTCTGTGGATGTGTGGGCAT |  |  |  |  |  |  |  |  |  |
|  |  | P: CGGGCCCTGTCACT |  |  |  |  |  |  |  |  |  |
|  | *ifn1* | F: CAGTATGCAGAGCGTGTGTCATT | 107.6 | 27.87 | 24.92 | 29.34 | 25.93 | 28.30 | 27.27 | 26.94 | 26.94 |
|  |  | R: TCTCCTCCCATCTGGTCCAG |  |  |  |  |  |  |  |  |  |
|  |  | P: CTGTGACTGGATCCGA |  |  |  |  |  |  |  |  |  |
|  | *il8* | F: TGGCCCTCCTGACCATTACT | 99.2 | 27.99 | 27.07 | 29.68 | 25.88 | 30.72 | 29.68 | 29.88 | 29.29 |
|  |  | R: GTCTCAATGCAGCGACATCG |  |  |  |  |  |  |  |  |  |
|  |  | P: ATGAGTCTGAGAGGCATG |  |  |  |  |  |  |  |  |  |
|  | *tapbp* | F: ACGGCAAGACTGACCGATTT | 103.5 | 26.14 | 27.19 | 27.32 | 25.86 | 28.39 | 26.91 | 27.22 | 29.10 |
|  |  | R: CACTTCAGCTTCCTGCAGGAT |  |  |  |  |  |  |  |  |  |
|  |  | P: AGCGGGGCTAGACTT |  |  |  |  |  |  |  |  |  |
| Endocrine disruption | *vtg1* | F: AAGTTCTGGGTAATGCTGGCC | 103 | 30.20 | - | - | - | 28.64 | - | - | 27.60 |
|  |  | R: CAAGACTGCATCAGCCTGGAC |  |  |  |  |  |  |  |  |  |
|  |  | P: TGCCCGTGTTTGGAA |  |  |  |  |  |  |  |  |  |
|  | *esr1* | F: GTGGAGGGTATGGCTGAGATCT | 103.5 | - | 28.53 | 31.35 | 29.78 | 30.28 | 31.76 | - | 33.98 |
|  |  | R: TCCTCAGGCTTCAGTTTAAGCAT |  |  |  |  |  |  |  |  |  |
|  |  | P: TGGCCACTGTGTCTC |  |  |  |  |  |  |  |  |  |
|  | *esr2a* | F: TTTGTGGACCTGTGCCTGTTC | 107.5 | 29.78 | 28.83 | 29.32 | 29.38 | - | 28.48 | - | - |
|  |  | R: CATGAGCCCTAGCATCAGCA |  |  |  |  |  |  |  |  |  |
|  |  | P: TTGGAGTGCTGCTGGT |  |  |  |  |  |  |  |  |  |
|  | *esr2b* | F: AACGAGGCCTGTCATTCCAG | 108.7 | 29.42 | 26.63 | 28.70 | 27.59 | 29.53 | 28.80 | - | 29.54 |
|  |  | R: TGTGTGAGAGCAGCATGAGGA |  |  |  |  |  |  |  |  |  |
|  |  | P: TCATCCCGCCTGGC |  |  |  |  |  |  |  |  |  |
|  | *ar* | F: AAGTGGTCAAGTGGGCCAAA | 106.4 | 28.69 | 28.75 | 28.26 | 27.27 | 29.06 | 27.33 | 30.13 | 30.13 |
|  |  | R: ACCCCCATCCATGAATGCT |  |  |  |  |  |  |  |  |  |
|  |  | P: TTGCCAGGTTTTCGGAAT |  |  |  |  |  |  |  |  |  |
|  | *cyp19a1b* | F: CGGACGACGTGAGACAGTGT | 127.5 | - | 29.43 | - | - | - | 29.81 | - | - |
|  |  | R: TCCTCATCTCCACCTCAGGG |  |  |  |  |  |  |  |  |  |
|  |  | P: TGATAGCAGCCCCAGACA |  |  |  |  |  |  |  |  |  |
| Circadian rhythm | *nr1d1* | F: TGTCCCGCGATGCTGTT | 110.5 | 25.34 | 26.22 | 27.85 | 23.12 | 26.46 | 24.18 | 26.10 | 26.58 |
|  |  | R: CCAGCATGCGCTGCTTCT |  |  |  |  |  |  |  |  |  |
|  |  | P: CTTTGGCAGAATACC |  |  |  |  |  |  |  |  |  |
|  | *nr1d2a* | F: TCGGATAGCGGCGATGA | 117.6 | 24.07 | 23.83 | 26.68 | 22.76 | 28.38 | 25.94 | - | - |
|  |  | R: GAATGTGCCCTGGTGACACA |  |  |  |  |  |  |  |  |  |
|  |  | P: AGGTTCCTCCTTCCAGG |  |  |  |  |  |  |  |  |  |
|  | *nr1d2b* | F: TCGGCATGTCCAAGGACTCT | 103.1 | 26.45 | 25.49 | 28.15 | 25.20 | 27.65 | 26.27 | 25.82 | 27.04 |
|  |  | R: CGCTGCTTCTCACGCTTTG |  |  |  |  |  |  |  |  |  |
|  |  | P: TGCGTTTCGGCCGCA |  |  |  |  |  |  |  |  |  |
|  | *arntl1a* | F: TCCAGCAGCCCGAGTAATG | 106.8 | 29.39 | 28.64 | 27.83 | 27.40 | 29.67 | 27.60 | 27.30 | 28.05 |
|  |  | R: GCCTCCAGAAGGCTCATGATC |  |  |  |  |  |  |  |  |  |
|  |  | P: CGAGGCAGCCATGG |  |  |  |  |  |  |  |  |  |
|  | *per1a/b* | F: CTGGACAACATTGCCTCAGAGTAC | 95.3 | 29.64 | 27.76 | 28.31 | 26.49 | 28.78 | 27.62 | - | - |
|  |  | R: GAAGGACACCGCCATGGA |  |  |  |  |  |  |  |  |  |
|  |  | P: CCCTCAAAAATACAGACACC |  |  |  |  |  |  |  |  |  |
|  | *per2* | F: GAGTACACCCTCAAAAACAATGACA | 100.2 | 30.98 | - | 32.04 | 28.61 | 31.15 | 27.72 | 29.97 | 30.91 |
|  |  | R: ATGTAGACGATCTTCCCCGTGAT |  |  |  |  |  |  |  |  |  |
|  |  | P: TGCGGTAGCGATATC |  |  |  |  |  |  |  |  |  |
|  | *cry1a/b* | F: CCGCCGGGACAAGGA | 103.1 | 25.90 | 25.28 | 29.82 | 24.91 | 26.34 | 27.90 | 26.28 | 25.86 |
|  |  | R: AGATGATTTCTACTCCGTGCTCTTC |  |  |  |  |  |  |  |  |  |
|  |  | P: TGGGCCGGTTAGC |  |  |  |  |  |  |  |  |  |
|  | *cry2* | F: TCCGCCTCTTCCTGAATGG | 102.9 | 26.87 | 26.57 | 27.79 | 26.33 | 27.44 | 26.48 | 26.60 | 27.02 |
|  |  | R: CCCACTACAGAGGCCTCAGTCT |  |  |  |  |  |  |  |  |  |
|  |  | P: TGGCATCTGTGCCC |  |  |  |  |  |  |  |  |  |
|  | *roraa/b* | F: TGCCGTGCCTTCGACTCT | 116 | 26.82 | 25.01 | 28.78 | 24.78 | 26.38 | 26.39 | 24.79 | 24.97 |
|  |  | R: CAGGCCCAGCATACTTTCCA |  |  |  |  |  |  |  |  |  |
|  |  | P: ACAACACAGTCTATTTCG |  |  |  |  |  |  |  |  |  |
|  | *clocka* | F: CGCAATGCAGCACCTGAA | 106.8 | 26.30 | 26.79 | 27.69 | 26.29 | 26.82 | 25.76 | 26.87 | 26.22 |
|  |  | R: ATGTTGGCCTCGATCATCCT |  |  |  |  |  |  |  |  |  |
|  |  | P: CTGGAGCAGAGGAC |  |  |  |  |  |  |  |  |  |
|  | *clockb* | F: CCTGGAGTCACTGGCCAAAT | 106 | 27.54 | 26.72 | 28.16 | 27.47 | 28.97 | 26.96 | 30.37 | 28.83 |
|  |  | R: CACGACTTCCCCTTCCCATA |  |  |  |  |  |  |  |  |  |
|  |  | P: CCACGAACACTTAATG |  |  |  |  |  |  |  |  |  |
|  | *npas2* | F: ACGCATGCTGCTCAACCA | 108 | 28.69 | 27.42 | - | 28.25 | 27.68 | 29.96 | - | - |
|  |  | R: CCTGGGAGCTAACACGGCTAT |  |  |  |  |  |  |  |  |  |
|  |  | P: CAGTGCAGACACTCGTA |  |  |  |  |  |  |  |  |  |
|  | *dbpa/b* | F: GAAGGGAGACCGTGCAAGTC | 111.8 | 26.28 | 26.51 | 28.45 | 25.68 | 30.04 | 30.42 | 31.97 | 35.23 |
|  |  | R: GGAGTCCTCATCCATGTCTGTGA |  |  |  |  |  |  |  |  |  |
|  |  | P: TGCGACGTTAAAGAC |  |  |  |  |  |  |  |  |  |
| Apoptosis | *casp3a/b* | F: AAAGGGATAGCTGCACAAGGG | 121.4 | 25.34 | 24.97 | 28.24 | 23.71 | 25.60 | 24.69 | - | - |
|  |  | R: TGTGTACATGTCAGCCGGAGG |  |  |  |  |  |  |  |  |  |
|  |  | P: CAGTCCCTCAGTGCAAA |  |  |  |  |  |  |  |  |  |
|  | *casp9* | F: GCCAGACAGTTGGTTCGAGAC | 102.8 | 28.45 | 26.59 | 29.88 | 27.40 | 27.92 | 27.72 | 28.23 | 28.02 |
|  |  | R: GGCTATGCTGCCCTTTCTCA |  |  |  |  |  |  |  |  |  |
|  |  | P: TCCCAGCTTTAATAGAG |  |  |  |  |  |  |  |  |  |
|  | *tp53* | F: CCTTTGAGGTGCGTGTGTGT | 118.6 | - | - | 25.79 | 23.95 | 25.28 | 24.24 | 24.79 | 24.37 |
|  |  | R: GGGTTGTCTCCTGCTGCTTCT |  |  |  |  |  |  |  |  |  |
|  |  | P: CTGGTCGAGACAGGAA |  |  |  |  |  |  |  |  |  |
|  | *pdcd10a/b* | F: AGGCTGAGAAGGAGAACCCAG | 104.8 | 25.42 | 23.96 | 28.89 | 24.24 | 25.91 | 26.17 | 26.44 | 26.14 |
|  |  | R: CACATCGTCTGCAGCCATTC |  |  |  |  |  |  |  |  |  |
|  |  | P: TGACCCAGGACATCAT |  |  |  |  |  |  |  |  |  |
|  | *chmp5a/b* | F: CCAAAGAGATGAAGGCGGC | 115 | 21.72 | 21.30 | 24.39 | 20.51 | 22.81 | 21.84 | 22.32 | 21.92 |
|  |  | R: TTGGCGTCCTCCATCATGT |  |  |  |  |  |  |  |  |  |
|  |  | P: AGGATCTCCAGGACCAG |  |  |  |  |  |  |  |  |  |
| Growth and metabolism | *ghr* | F: TAACCGGGAGCCACTTTGAC | 101.9 | 27.61 | 27.81 | 28.65 | 26.83 | 27.89 | 27.05 | 28.18 | 27.09 |
|  |  | R: ATTGACCTCACGGTACTGCACC |  |  |  |  |  |  |  |  |  |
|  |  | P: TGAGCTGGGAGCCG |  |  |  |  |  |  |  |  |  |
|  | *igf1* | F: TTCAAGAGTGCGATGTGCTGT | 108.5 | 28.14 | 25.93 | 29.22 | 26.78 | 26.63 | 27.83 | 29.76 | 28.72 |
|  |  | R: CGCCGAAGTCAGGGTTAGG |  |  |  |  |  |  |  |  |  |
|  |  | P: ACACCCTCTCACTGCT |  |  |  |  |  |  |  |  |  |
|  | *igf2* | F: ATGTGGAGGAGAACTGGTGGAC | 101.6 | 27.57 | 26.94 | 27.91 | 26.17 | 28.15 | 26.70 | 28.82 | 28.51 |
|  |  | R: CCTGCTGGTTGGCCTACTGA |  |  |  |  |  |  |  |  |  |
|  |  | P: TGCAGTTCGTCTGTGAAG |  |  |  |  |  |  |  |  |  |
|  | *igfbp1* | F: AACTGTGCGGAATCTACACGG | 111.9 | - | 30.39 | - | 29.25 | 31.08 | 29.45 | - | - |
|  |  | R: CTGGCTGCGAATAAGGGAGT |  |  |  |  |  |  |  |  |  |
|  |  | P: TGCACCCCAATACC |  |  |  |  |  |  |  |  |  |
|  | *igfbp2* | F: AGCTGCATGTCCTAAGCTTGC | 93 | - | - | 32.48 | 31.69 | 30.53 | - | - | - |
|  |  | R: GCCTTCTAACCGGGAACACA |  |  |  |  |  |  |  |  |  |
|  |  | P: TCAGGGAGCCCGGCT |  |  |  |  |  |  |  |  |  |
|  | *ampka1* | F: AAGTTTGAGTGCACCGAGGAG | 106.2 | 26.55 | 25.92 | 28.80 | 25.34 | 27.66 | 26.07 | 25.50 | 25.56 |
|  |  | R: GACATAATGCGGCGGTTGTC |  |  |  |  |  |  |  |  |  |
|  |  | P: CGCAACCACCACGAC |  |  |  |  |  |  |  |  |  |
|  | *ldhb* | F: CCCCCAACTGCACCCTTATT | 102 | - | 22.58 | 25.37 | 22.96 | 22.87 | 23.18 | 24.27 | 25.00 |
|  |  | R: AATCCGCTCAACTTCCACGT |  |  |  |  |  |  |  |  |  |
|  |  | P: CCAACCCAGTGGACGT |  |  |  |  |  |  |  |  |  |
|  | *pck1* | F: ATATGAGAACTGCTGGCTGGC | 125.2 | - | - | - | 29.86 | 30.35 | - | 29.90 | 30.22 |
|  |  | R: CACCTACTCGTGGAGACGGAA |  |  |  |  |  |  |  |  |  |
|  |  | P: CCCAGAGACGTGGCC |  |  |  |  |  |  |  |  |  |
|  | *fasn* | F: TGTGGGAGGTGTAGTCAAGCC | 109.5 | 24.53 | 23.49 | 26.59 | 24.48 | 26.17 | 24.62 | 24.66 | 25.49 |
|  |  | R: TCCCTGGGCCATGTATCTGA |  |  |  |  |  |  |  |  |  |
|  |  | P: AGGTGGAGGAGGCC |  |  |  |  |  |  |  |  |  |
|  | *cpt1a* | F: TACAGCTGGCCCAATTCAGG | 112.7 | 26.80 | 26.22 | 27.84 | 25.56 | 27.25 | 27.29 | 29.30 | 26.64 |
|  |  | R: AACCGTCTCTGTCCTACCCTCA |  |  |  |  |  |  |  |  |  |
|  |  | P: TCAATGACCCGGATGTT |  |  |  |  |  |  |  |  |  |
|  | *cpt1b* | F: GGACCAGTCCTGATGCCTTC | 119.5 | 26.69 | 26.45 | 29.08 | 26.24 | 28.21 | 30.14 | 29.43 | 26.72 |
|  |  | R: TCGAGGCCTCATACGTCAGAC |  |  |  |  |  |  |  |  |  |
|  |  | P: ACTGCAGCTGGCTCA |  |  |  |  |  |  |  |  |  |
|  | *cs* | F: TTGATTGCCAAGTTGCCGT | 121.6 | 23.51 | 23.24 | 25.17 | 21.92 | 23.13 | 21.83 | 22.98 | 23.19 |
|  |  | R: ATGTTGGCGAAGTTAGCGGA |  |  |  |  |  |  |  |  |  |
|  |  | P: CAGCAGCATTGGC |  |  |  |  |  |  |  |  |  |
|  | *ldha* | F: CGTCAAGTACAGCCCCAACG | 104.7 | 28.25 | 28.51 | 29.63 | 28.09 | 27.69 | 28.94 | 29.36 | 28.73 |
|  |  | R: GCCACGTAGGTCAGGATGTCA |  |  |  |  |  |  |  |  |  |
|  |  | P: TGCTGGTCGTCTCCAA |  |  |  |  |  |  |  |  |  |
|  | *lipea/b* | F: ACACTGGTCAAGGTGTTGCAGT | 111.6 | 26.98 | 27.20 | 28.62 | 27.04 | 27.78 | 25.63 | 28.70 | 26.97 |
|  |  | R: TGCAGTTAGCGGCAATGTAGC |  |  |  |  |  |  |  |  |  |
|  |  | P: CTTCTGCACATCATCCA |  |  |  |  |  |  |  |  |  |
|  | *lpl* | F: TGACAGCGCTGTACAAGAGGG | 113.3 | 26.76 | 26.59 | 28.39 | 27.19 | 27.10 | 26.66 | 28.89 | 27.76 |
|  |  | R: GGAGGTGAGGTAGTGCTGCTG |  |  |  |  |  |  |  |  |  |
|  |  | P: TGGACTGGCTGACACGG |  |  |  |  |  |  |  |  |  |
|  | *ctsd* | F: TTCACAGACATCGCCTGCTT | 119.4 | 23.00 | 22.69 | 25.69 | 23.46 | 23.52 | 23.40 | 22.00 | 23.71 |
|  |  | R: GGTACCCAGACAGACTGCCAG |  |  |  |  |  |  |  |  |  |
|  |  | P: CCACAAGTATAACGGTGCC |  |  |  |  |  |  |  |  |  |
| Detoxification | *cyp1a* | F: TACAGCTGGCCCAATTCAGG | 98.7 | 26.57 | 27.40 | 27.66 | 24.79 | 26.56 | 25.44 | 28.16 | 26.40 |
|  |  | R: AACCGTCTCTGTCCTACCCTCA |  |  |  |  |  |  |  |  |  |
|  |  | P: TCAATGACCCGGATGTT |  |  |  |  |  |  |  |  |  |
|  | *gstp1* | F: GGTGACAAGCCTTCGTTTGC | 112.5 | 23.59 | 22.57 | 26.83 | 22.12 | 23.85 | 23.90 | 22.78 | 22.89 |
|  |  | R: CAAAGCTCTTCAGGGAGGGG |  |  |  |  |  |  |  |  |  |
|  |  | P: TGAAGTGCTGCTCAAC |  |  |  |  |  |  |  |  |  |
|  | *gpx1a* | F: CCACCCCTTGTTTGTGTATCTCA | 119.7 | - | 24.10 | 26.71 | 23.66 | 24.91 | 24.20 | - | - |
|  |  | R: GGGCTCCACATGATGAACTTG |  |  |  |  |  |  |  |  |  |
|  |  | P: CCATTCCCCTCCGATG |  |  |  |  |  |  |  |  |  |
|  | *cat* | F: TGGGCCGCTACAACAGTACTG | 103.8 | 25.66 | 25.14 | 28.06 | 26.51 | 24.93 | 26.47 | 26.41 | 25.81 |
|  |  | R: CTCGTTCAGCACCTTAGTGAAGAA |  |  |  |  |  |  |  |  |  |
|  |  | P: CGTCACACAGGTGCGTA |  |  |  |  |  |  |  |  |  |
|  | *glul* | F: TGAAGTCATGCCTGCACAGTG | 120.7 | 25.24 | 23.00 | 27.17 | 24.12 | 27.39 | 25.51 | 24.61 | 25.01 |
|  |  | R: CACACCCGGTGGAGAATGA |  |  |  |  |  |  |  |  |  |
|  |  | P: CAGGTTGGCCCTTGT |  |  |  |  |  |  |  |  |  |
|  | *sod2* | F: GATGGCTGGGCTTTGACAA | 117.4 | 25.65 | 22.75 | 27.51 | 23.81 | 25.03 | 24.69 | 25.13 | 24.71 |
|  |  | R: CCTGCAGTGGGTCTTGATTAGG |  |  |  |  |  |  |  |  |  |
|  |  | P: AAGCTCCGTATCACAGC |  |  |  |  |  |  |  |  |  |
|  | *sod1* | F: CAACACCAACGGCTGTATGAGT | 93.5 | 31.82 | 21.85 | 24.61 | 25.28 | 24.37 | 24.07 | 23.82 | 22.85 |
|  |  | R: CCTCCGTGGGTCTTGTTGTG |  |  |  |  |  |  |  |  |  |
|  |  | P: CGGACCCCACTTCA |  |  |  |  |  |  |  |  |  |
|  | *nfe2l2a* | F: GGTGGCTACAGCGATTCAGAC | 123 | 24.79 | 23.77 | 27.17 | 23.86 | 25.17 | 24.80 | 24.30 | - |
|  |  | R: GAAAGAGGTGTCTGCAGGCC |  |  |  |  |  |  |  |  |  |
|  |  | P: AGTAACCCTGGGAGTGC |  |  |  |  |  |  |  |  |  |
| Hypoxia | *hif1a* | F: GCTGTGGGCTGAAGAGTGATC | 117.6 | 26.73 | 26.28 | 28.37 | 26.45 | 27.26 | 26.78 | 24.78 | 25.93 |
|  |  | R: CTGGGTTCTCCTTCAGCTGG |  |  |  |  |  |  |  |  |  |
|  |  | P: TCTCTGAGGCCTCCGAG |  |  |  |  |  |  |  |  |  |
|  | *epor* | F: TAAAGTGGCTCTGCTGCTCCA | 109 | 25.69 | 26.85 | 27.43 | 26.15 | 27.43 | 26.72 | - | 31.45 |
|  |  | R: CCAGAAGCAGGTGAGGTCTGA |  |  |  |  |  |  |  |  |  |
|  |  | P: CTGAGCCAGAGAATC |  |  |  |  |  |  |  |  |  |
|  | *vegfc* | F: AACCACACCGTGTGCAATTG | 93.1 | 27.10 | 26.82 | 27.71 | 25.39 | 26.86 | 26.21 | - | - |
|  |  | R: TGTGGTTCTTTGGGCAGGTT |  |  |  |  |  |  |  |  |  |
|  |  | P: ACTGGACGCCTACAGAC |  |  |  |  |  |  |  |  |  |
|  | *slc2a1a* | F: TGAGCATCGTGGCCATCTT | 125.8 | 23.41 | 23.17 | 25.37 | 22.74 | 24.37 | 22.50 | 22.56 | 23.86 |
|  |  | R: AACAGCTCAGCCACGATGAA |  |  |  |  |  |  |  |  |  |
|  |  | P: TTGGCCCGGGTCC |  |  |  |  |  |  |  |  |  |
|  | *mb* | F: ATGGTTCTGAAGTGCTGGGG | 125.1 | 28.11 | - | 32.35 | 28.87 | 30.05 | 29.54 | 26.91 | 28.42 |
|  |  | R: AAACAGACGGCTCAGAACCAG |  |  |  |  |  |  |  |  |  |
|  |  | P: AGGCTGACTACAACAAA |  |  |  |  |  |  |  |  |  |
|  | *hk1* | F: ATGCTGGAGGACATTCGCA | 118.3 | 26.47 | 25.96 | - | 27.57 | - | 27.36 | 26.40 | 27.67 |
|  |  | R: TTCGCCCATGTACATCCCA |  |  |  |  |  |  |  |  |  |
|  |  | P: AGGGGCTCTCTAAAC |  |  |  |  |  |  |  |  |  |
|  | *pfkm* | F: TGGTATCTACACCGGAGCCAA | 114.8 | 30.75 | 29.20 | - | 28.56 | 29.29 | 28.75 | 30.02 | 28.78 |
|  |  | R: TGCAGCATCATGGACACACTC |  |  |  |  |  |  |  |  |  |
|  |  | P: CCAGGGTCTGGTGGAT |  |  |  |  |  |  |  |  |  |
|  | *aldoaa* | F: CAACGGAGAGACCACCACTCA | 107.9 | 23.02 | 23.55 | 24.20 | 22.32 | 22.99 | 21.59 | 22.58 | 23.27 |
|  |  | R: ACGCCACTTAGCAAAGTCAGC |  |  |  |  |  |  |  |  |  |
|  |  | P: TGTACGAGCGGTGTGC |  |  |  |  |  |  |  |  |  |
|  | *eno1a* | F: TCCCTGCCTTCAACGTGATC | 100.6 | 22.93 | 22.39 | 24.29 | 21.19 | 22.90 | 22.01 | 23.05 | 23.30 |
|  |  | R: CCTCATGGCCTCCTTGAAGG |  |  |  |  |  |  |  |  |  |
|  |  | P: CGGAGGTTCCCACGCA |  |  |  |  |  |  |  |  |  |
|  | *pgk1* | F: AGATGATCATCGGTGGTGGC | 104.3 | 24.91 | 24.40 | 25.46 | 24.19 | 25.45 | 23.69 | 25.28 | 25.74 |
|  |  | R: CATACAGGGAGGTGCCGATC |  |  |  |  |  |  |  |  |  |
|  |  | P: CCTTCACCTTCCTCAATG |  |  |  |  |  |  |  |  |  |
| Endogenous control | *rpl7* | F: TCGTCATCAGGATCAGGGGT | 118.4 | 18.60 | 17.99 | 20.53 | 18.45 | 19.27 | 18.89 | 18.76 | 19.43 |
|  |  | R: GAAGATCTGACGCAGACGCA |  |  |  |  |  |  |  |  |  |
|  |  | P: CAAGGTGCGCAAGGT |  |  |  |  |  |  |  |  |  |
|  | *rps9* | F: TTCTCCCTGCGTTCACCATAC | 116.2 | 20.22 | 19.61 | 21.48 | 18.86 | 21.14 | 20.06 | 19.59 | 20.27 |
|  |  | R: GGCCCTTCTTGGCATTCTTT |  |  |  |  |  |  |  |  |  |
|  |  | P: CCCGGCCGTGTCAA |  |  |  |  |  |  |  |  |  |
|  | *ef1a* | F: GAAGCTTGAGGACAACCCCA | 117.3 | 22.54 | 22.73 | 24.56 | 22.23 | 23.26 | 22.92 | 22.78 | 23.68 |
|  |  | R: GAAGCTCTCCACACACATGGG |  |  |  |  |  |  |  |  |  |
|  |  | P: CCGCCATCATCGTCAT |  |  |  |  |  |  |  |  |  |
|  | *rpl13a* | F: CACTGGAGAGGCTGAAGGTGT | 116.7 | 20.53 | 20.67 | 21.76 | 19.98 | 21.30 | 19.28 | 19.93 | 21.27 |
|  |  | R: GTGGGCTTCAGACGGACAAT |  |  |  |  |  |  |  |  |  |
|  |  | P: CATGGTCGTACCTGCT |  |  |  |  |  |  |  |  |  |

Supplementary Table 2: TaqMan primer and probe sequences, PCR efficiencies, and mean C_T_ levels for the 112 candidate genes for liver tissue across eight salmonid species

| Biological function | Gene symbol | Primer and probe sequences | Efficiency (%) | C_T_ level | | | | | | | |
| --- | --- | --- | --- | --- | --- | --- | --- | --- | --- | --- | --- |
|  |  |  |  | *S.*  *salar* | *S. trutta* | *O.*  *mykiss* | *O. tshawytscha* | *S.*  *fontinalis* | *S.*  *alpinus* | *C.*  *clupeaformis* | *C.*  *hoyi* |
| Stress | *gr1* | F: GTCTTTGGCCTGTATCCCCC | 108.4 | 24.95 | 24.49 | 25.87 | 24.42 | - | - | 27.28 | 26.60 |
|  |  | R: AGCTCGACATCCCTGATCCA |  |  |  |  |  |  |  |  |  |
|  |  | P: TGCCCTCGGTCAGTGA |  |  |  |  |  |  |  |  |  |
|  | *gr2* | F: ATGGCAGACCAGTGTGAACAGAT | 114 | 27.57 | 28.01 | 27.40 | 26.42 | 25.89 | 27.30 | 28.12 | 25.64 |
|  |  | R: GAGCAGCAGCAGAACCTTCAT |  |  |  |  |  |  |  |  |  |
|  |  | P: AGACTGCAGGTGTCTCA |  |  |  |  |  |  |  |  |  |
|  | *mr* | F: CATAGTCAATGTCAGCTGCTCCC | 103.1 | 26.00 | 25.40 | 25.57 | 24.31 | 25.68 | 26.59 | 26.02 | 25.10 |
|  |  | R: GCTGCTGCTCTGGCTTCTTCT |  |  |  |  |  |  |  |  |  |
|  |  | P: CCAGCAGCAACACA |  |  |  |  |  |  |  |  |  |
|  | *hsd11b2* | F: CTGTCTAGCAGCGTACGGAGC | 106.5 | 29.36 | 28.94 | 26.90 | 26.78 | 29.43 | 28.15 | 29.74 | 28.32 |
|  |  | R: ATGGTGGACACTTTGACCCC |  |  |  |  |  |  |  |  |  |
|  |  | P: CCTGTTCATCAACACACT |  |  |  |  |  |  |  |  |  |
|  | *hsp70a* | F: GAGAACACTGTCCTCCAGCTCC | 108.5 | 27.63 | 27.39 | 27.22 | 25.84 | 26.18 | 29.23 | 26.70 | 30.33 |
|  |  | R: CCCTGAAGAGGTCGGAACAC |  |  |  |  |  |  |  |  |  |
|  |  | P: ACACCTCCATCACCAGG |  |  |  |  |  |  |  |  |  |
|  | *hsp90aa* | F: AAGATCGAGGTCACCCCTGA | 92.2 | 30.67 | - | - | - | - | 32.57 | - | - |
|  |  | R: GGTGCCAGACTTTGCAATGG |  |  |  |  |  |  |  |  |  |
|  |  | P: CGGCATCGGCATGA |  |  |  |  |  |  |  |  |  |
|  | *hsp90ba* | F: GAGGTGGAGGAGGACGAGTACA | 108.8 | 23.13 | 24.47 | 26.71 | 23.54 | 23.54 | 25.91 | 23.25 | 25.55 |
|  |  | R: GCTGTGAAGTGGATGTGGGA |  |  |  |  |  |  |  |  |  |
|  |  | P: TCTCCAGGGACACAGAC |  |  |  |  |  |  |  |  |  |
|  | *hsf1* | F: CCCAAGTTCAGCAGGCAGTAC | 92.4 | 24.96 | 24.18 | 24.84 | 24.41 | 24.30 | 24.42 | 24.94 | 25.11 |
|  |  | R: GCCGTGAAGAGACCGGTACT |  |  |  |  |  |  |  |  |  |
|  |  | P: TGCAGGGCTCGCCT |  |  |  |  |  |  |  |  |  |
|  | *cirbpa* | F: CGGGAAGGTCTCGTGGATT | 116.9 | 22.84 | 21.84 | 23.67 | 22.34 | 21.29 | 23.03 | 22.09 | 20.61 |
|  |  | R: TGGTTCTGCCATCGACAGACT |  |  |  |  |  |  |  |  |  |
|  |  | P: CATTGGAGGGAATGAA |  |  |  |  |  |  |  |  |  |
|  | *crfb1* | F: TCATTGCTTTCTTACCGCGC | 106.4 | - | - | - | - | 29.55 | - | - | - |
|  |  | R: AGGAGGGGAAGAGACTGTTGCT |  |  |  |  |  |  |  |  |  |
|  |  | P: CCGCTCCACATCACGA |  |  |  |  |  |  |  |  |  |
|  | *avt* | F: CTAGACCCAGACTGCCTAGAGGAC | 94 | 30.83 | 30.00 | 31.69 | 31.04 | 29.60 | 30.32 | - | - |
|  |  | R: GCCAAACCACCCATTAAGGC |  |  |  |  |  |  |  |  |  |
|  |  | P: ACGTCAGTCACCCAGCGA |  |  |  |  |  |  |  |  |  |
|  | *mmp2* | F: CGCTGTGGAGTTCCTGATGTT | 102.7 | 24.13 | 23.81 | 26.10 | 23.91 | 25.30 | 28.55 | 30.14 | 29.55 |
|  |  | R: AGGTCAGGAGAGTGGCCTAGAA |  |  |  |  |  |  |  |  |  |
|  |  | P: AGGAAACCCAAGTGGCA |  |  |  |  |  |  |  |  |  |
|  | *mmp9* | F: ACGGCAAAGCATGTGTGTTC | 115.1 | 27.61 | 28.72 | 29.58 | - | 27.34 | 27.50 | - | - |
|  |  | R: GGAACACCAGCGGTATCCAT |  |  |  |  |  |  |  |  |  |
|  |  | P: AGGTTGCACGACGGAA |  |  |  |  |  |  |  |  |  |
|  | *mt* | F: GCTCAAAAACTGGACGCTGC | 98.7 | 28.36 | - | - | 30.08 | 28.23 | - | - | - |
|  |  | R: GGCAGCAGGAACAACAACTTT |  |  |  |  |  |  |  |  |  |
|  |  | P: TACAAACTGCGGATGTGC |  |  |  |  |  |  |  |  |  |
|  | *mtA* | F: TGGATCCTTGTGAATGCTCCA | 101.2 | 24.55 | 22.02 | 23.04 | 26.47 | 19.43 | 20.80 | 22.19 | 21.20 |
|  |  | R: CTTACAACTGGTGCATGCGC |  |  |  |  |  |  |  |  |  |
|  |  | P: CGGTGGATCCTGCAAG |  |  |  |  |  |  |  |  |  |
|  | *mtB* | F: GAAAAGTTGCTGCCCCTGC | 112.6 | 24.04 | 20.17 | 23.54 | 25.53 | 22.96 | 22.65 | 25.60 | 22.66 |
|  |  | R: AACAGCTGGTATCGCAGGTCTT |  |  |  |  |  |  |  |  |  |
|  |  | P: TTCAGGCTGTGTGTGCA |  |  |  |  |  |  |  |  |  |
|  | *hsd20b2* | F: ACATTGTACTGGTCAGCCGGT | 109.7 | 28.82 | 28.60 | - | 28.85 | 25.33 | 27.25 | 27.21 | 26.70 |
|  |  | R: TGGCCCTCTGTGAAGTCTGTC |  |  |  |  |  |  |  |  |  |
|  |  | P: CCAACATGGACGCCAGA |  |  |  |  |  |  |  |  |  |
|  | *hsp30* | F: TGCTGTGTTCCCGAGGATTC | 123.5 | - | 33.50 | 27.96 | - | - | - | - | - |
|  |  | R: TCTCTGCAGTAGATCCCGCTG |  |  |  |  |  |  |  |  |  |
|  |  | P: TGTGCGCAGTCTAT |  |  |  |  |  |  |  |  |  |
|  | *serpinh1a* | F: AGCGCTGTGAAGTCCATCAA | 96.3 | 27.74 | 26.03 | 26.96 | 28.00 | 24.70 | 26.71 | 30.09 | 28.81 |
|  |  | R: TGATGATCATGGCCCCATC |  |  |  |  |  |  |  |  |  |
|  |  | P: CGGCCAAGTCCACCGA |  |  |  |  |  |  |  |  |  |
|  | *hspa4* | F: ACTGCTGAGACCGCAATGAA | 116.4 | 25.24 | 24.04 | 28.42 | 24.32 | 23.16 | 26.61 | 25.69 | 25.86 |
|  |  | R: TGCGTCAGTGTAGAAGCTGGG |  |  |  |  |  |  |  |  |  |
|  |  | P: ACCTGTGGCTGATTGT |  |  |  |  |  |  |  |  |  |
|  | *hsp7c* | F: TAGCCAACGACCAGGGAAAC | 106.8 | 25.15 | 24.77 | 27.03 | 23.15 | 23.03 | 25.73 | 30.19 | 30.97 |
|  |  | R: CGTCCCCAATCAGCCTTTCT |  |  |  |  |  |  |  |  |  |
|  |  | P: ACCCAGTTATGTCGCCTT |  |  |  |  |  |  |  |  |  |
|  | *pomc-a1* | F: AGAACAGCATCCTGGAGTGCA | 89.4 | - | - | 30.50 | - | - | - | - | - |
|  |  | R: GGGAGTTGGGTTGGAGATGG |  |  |  |  |  |  |  |  |  |
|  |  | P: ACCTCACCGCCGAAT |  |  |  |  |  |  |  |  |  |
| Neural plasticity | *bdnf* | F: TCCTTCCTCTGTGGTCACCAC | - | - | - | - | - | - | - | - | - |
|  |  | R: AGGCACTTGGTTGCTGATCA |  |  |  |  |  |  |  |  |  |
|  |  | P: TGTCGACCTGTATGCAT |  |  |  |  |  |  |  |  |  |
|  | *pcna* | F: TGGAGGCTCTGAAGGACCTG | 107.8 | 26.20 | 24.50 | 27.40 | 25.53 | 23.20 | 26.23 | 26.22 | 26.26 |
|  |  | R: GAGCTGAACTAGGGAGACGTGG |  |  |  |  |  |  |  |  |  |
|  |  | P: CTGTTGGGACGTGAGCT |  |  |  |  |  |  |  |  |  |
|  | *neurod1* | F: ATGTCATCAGACCCCCAGGA | 98.6 | - | - | - | 31.37 | 28.91 | - | - | - |
|  |  | R: GATTCGAGTGCGCCGTTTT |  |  |  |  |  |  |  |  |  |
|  |  | P: CGACCTGGACTGATGA |  |  |  |  |  |  |  |  |  |
|  | *dcx* | F: GTCTGTGAACGTGAAGGCCTC | 100 | 29.36 | 25.26 | 29.98 | 31.30 | 29.99 | 30.13 | 30.93 | 29.92 |
|  |  | R: TTGGGCCGAACAAACTCCT |  |  |  |  |  |  |  |  |  |
|  |  | P: AGCCAGAAGAACCTG |  |  |  |  |  |  |  |  |  |
|  | *neurog1* | F: AGAGCGAGGAATGAAGCCAC | - | - | - | - | - | - | - | - | - |
|  |  | R: GTTTCTCTCGCGGTCGTTTG |  |  |  |  |  |  |  |  |  |
|  |  | P: CGTGGTCAAGAAGAA |  |  |  |  |  |  |  |  |  |
|  | *sox2* | F: TCAGACAGGAGACCTACGGGAC | 89.9 | - | - | - | 28.29 | 28.60 | - | - | 30.52 |
|  |  | R: TTGGGACATGTGGAGTCTGC |  |  |  |  |  |  |  |  |  |
|  |  | P: TTACCGGGCGCTGAG |  |  |  |  |  |  |  |  |  |
| Appetite regulation | *agrp* | F: ATACTGCTGGACCCTGTGCCT | - | - | - | - | - | - | - | - | - |
|  |  | R: CGGGATGGGAGTCGTCTAGA |  |  |  |  |  |  |  |  |  |
|  |  | P: CCTGATCCAGCTGGCTA |  |  |  |  |  |  |  |  |  |
|  | *cart* | F: TGACCAACGAAAAGCAACTGC | 95.7 | - | 29.30 | - | - | - | - | - | - |
|  |  | R: AGGGACTTGGCCGAATTTCT |  |  |  |  |  |  |  |  |  |
|  |  | P: CTGCAGACCAAGAGA |  |  |  |  |  |  |  |  |  |
|  | *cck-I* | F: GGGTCCCAGCCACAAGATAA | 96.2 | 31.06 | 30.94 | - | 29.05 | 29.83 | - | - | - |
|  |  | R: CTCGTACTCCTCTGCACTGCG |  |  |  |  |  |  |  |  |  |
|  |  | P: TGGGCTGGATGGAC |  |  |  |  |  |  |  |  |  |
|  | *lepr* | F: CGCTGTATGCACATCAACGG | 99.9 | 32.05 | 30.79 | 31.30 | 30.87 | 30.62 | - | 33.34 | 33.28 |
|  |  | R: GTCAGGAGCTCTGCTGTTGTGA |  |  |  |  |  |  |  |  |  |
|  |  | P: CCGGGACCTGGAGTGA |  |  |  |  |  |  |  |  |  |
|  | *lepa* | F: AAGCCTGCCTTCCATAGTGGA | 116.2 | 31.33 | 29.19 | 29.72 | 31.40 | 28.18 | 32.02 | 29.14 | 29.49 |
|  |  | R: TGAGGTCTGCCCAGTCTAGGA |  |  |  |  |  |  |  |  |  |
|  |  | P: TGGGATTCTACCAGGACC |  |  |  |  |  |  |  |  |  |
|  | *npy* | F: AACCTCATCACAAGGCAGAGGT | 113 | 29.99 | - | - | - | 27.54 | 29.70 | - | - |
|  |  | R: ACGTGTCTGTGCTCTCCTTCAG |  |  |  |  |  |  |  |  |  |
|  |  | P: AGGTCCAGCCCTGAC |  |  |  |  |  |  |  |  |  |
| Osmoregulation | *cftr* | F: CAGGCAGAAAAGAGCAGGGA | 90.3 | - | - | - | - | 25.61 | - | - | - |
|  |  | R: TCTCCATCACCTCCTCCCAG |  |  |  |  |  |  |  |  |  |
|  |  | P: TCGTCGCCTGGCTC |  |  |  |  |  |  |  |  |  |
|  | *atpa1a* | F: GGTGGTTGGAGATCTGGTGG | - | - | - | - | - | - | - | - | - |
|  |  | R: CACCAGTGAGGGAGGAGTTGTC |  |  |  |  |  |  |  |  |  |
|  |  | P: ATTTGCGTATTGTCTCTG |  |  |  |  |  |  |  |  |  |
|  | *atpa1b* | F: GGATGCTTGGCTGGGATTT | 98.5 | 22.53 | 22.23 | 24.40 | 22.57 | 21.51 | 25.01 | 23.24 | 22.91 |
|  |  | R: CTCTGTCCTGGTAGGTGGGTTT |  |  |  |  |  |  |  |  |  |
|  |  | P: CAGGCCCTGCTGCT |  |  |  |  |  |  |  |  |  |
|  | *crhbp* | F: GAGCCCAACCAGGTCATCAA | 84.7 | - | 30.13 | - | 29.53 | 30.76 | - | 32.02 | - |
|  |  | R: CTCTCCCTTCATCACCCAGC |  |  |  |  |  |  |  |  |  |
|  |  | P: CGTTGACATCGACTGCA |  |  |  |  |  |  |  |  |  |
| Immune function | *cam* | F: GAGGAGCAGATTGCCGAGTT | 110 | 24.09 | 22.71 | 25.42 | 22.79 | 21.42 | 24.89 | 23.61 | 22.12 |
|  |  | R: CATGACAGTGCCCAGCTCTTT |  |  |  |  |  |  |  |  |  |
|  |  | P: CGCTCTTTGACAAGGA |  |  |  |  |  |  |  |  |  |
|  | *mhc-I* | F: AGTCCCTCCCTCAGTGTCTCTG | 106 | 23.18 | 22.97 | 23.82 | 20.95 | 20.86 | 22.59 | 22.91 | 23.18 |
|  |  | R: AGGACACCATGACTCCACTGG |  |  |  |  |  |  |  |  |  |
|  |  | P: AGTGACCTGCCACGCG |  |  |  |  |  |  |  |  |  |
|  | *mhc-II* | F: CTCACAGCAGCATCTACCCCA | 111.3 | 34.24 | 31.07 | 26.61 | 25.30 | 27.12 | - | 26.68 | 26.46 |
|  |  | R: CACCTGACTCTGACAGGTGCAG |  |  |  |  |  |  |  |  |  |
|  |  | P: TGGAGAACACCCTCATCT |  |  |  |  |  |  |  |  |  |
|  | *prdx* | F: GGATCAACACCCCCAGGAA | 103.7 | 25.61 | 23.60 | 25.34 | 24.59 | 23.20 | 24.57 | 25.23 | 24.30 |
|  |  | R: GTCCTCCTTCAGCACTCCGTA |  |  |  |  |  |  |  |  |  |
|  |  | P: CCCTTGTGGCTGACCT |  |  |  |  |  |  |  |  |  |
|  | *il-1β* | F: TCAGGGTCTGGATCTGGAGG | 124.3 | 28.63 | - | - | 27.81 | 28.67 | - | - | - |
|  |  | R: CTCGGTTCCCATGGTAACCC |  |  |  |  |  |  |  |  |  |
|  |  | P: ACCCCATCACCATGC |  |  |  |  |  |  |  |  |  |
|  | *ifnγ* | F: TGTTTTCCCCAAGGACACGT | 122.6 | 29.32 | 30.11 | - | 30.37 | 29.23 | - | - | - |
|  |  | R: CCGATACACGTCCAGAACCA |  |  |  |  |  |  |  |  |  |
|  |  | P: TGAGCGGAGGGTGTT |  |  |  |  |  |  |  |  |  |
|  | *tnfα* | F: CTCAACTCTGTACGCACCGTG | 96 | 30.17 | 29.42 | 28.92 | 29.87 | 29.11 | - | - | - |
|  |  | R: TCTCCCTTCTCCAGGCTGAA |  |  |  |  |  |  |  |  |  |
|  |  | P: CGAGGCTGCGAGTGA |  |  |  |  |  |  |  |  |  |
|  | *mx* | F: GGAGGAGATTGAGGACCCCT | 98.3 | 29.19 | 27.96 | 29.34 | 26.51 | 28.23 | 28.55 | 28.10 | 28.61 |
|  |  | R: ATCACTGATACCCACCCCCA |  |  |  |  |  |  |  |  |  |
|  |  | P: TGAAGCCCAGGATGAA |  |  |  |  |  |  |  |  |  |
|  | *stat1* | F: CAGAAAGGCTTCCTGGAGGG | 110.7 | 25.12 | 24.36 | 27.08 | 24.98 | 24.44 | 26.95 | 26.31 | 25.08 |
|  |  | R: CTCTGTGGATGTGTGGGCAT |  |  |  |  |  |  |  |  |  |
|  |  | P: CGGGCCCTGTCACT |  |  |  |  |  |  |  |  |  |
|  | *ifn1* | F: CAGTATGCAGAGCGTGTGTCATT | 116.1 | 29.67 | 27.70 | 29.77 | 28.81 | 28.52 | - | 29.45 | 28.24 |
|  |  | R: TCTCCTCCCATCTGGTCCAG |  |  |  |  |  |  |  |  |  |
|  |  | P: CTGTGACTGGATCCGA |  |  |  |  |  |  |  |  |  |
|  | *il8* | F: TGGCCCTCCTGACCATTACT | 107.4 | 29.58 | 30.51 | 31.52 | 30.44 | 31.69 | 31.12 | 29.57 | 29.40 |
|  |  | R: GTCTCAATGCAGCGACATCG |  |  |  |  |  |  |  |  |  |
|  |  | P: ATGAGTCTGAGAGGCATG |  |  |  |  |  |  |  |  |  |
|  | *tapbp* | F: ACGGCAAGACTGACCGATTT | 103.8 | 28.61 | 28.41 | 28.66 | 28.59 | 27.81 | 27.92 | 29.69 | 31.85 |
|  |  | R: CACTTCAGCTTCCTGCAGGAT |  |  |  |  |  |  |  |  |  |
|  |  | P: AGCGGGGCTAGACTT |  |  |  |  |  |  |  |  |  |
| Endocrine disruption | *vtg1* | F: AAGTTCTGGGTAATGCTGGCC | 116 | 29.93 | 28.88 | - | 29.22 | 28.91 | 27.32 | 22.62 | 16.03 |
|  |  | R: CAAGACTGCATCAGCCTGGAC |  |  |  |  |  |  |  |  |  |
|  |  | P: TGCCCGTGTTTGGAA |  |  |  |  |  |  |  |  |  |
|  | *esr1* | F: GTGGAGGGTATGGCTGAGATCT | 117 | 31.43 | 23.06 | 26.62 | 23.74 | 26.99 | 25.19 | 34.28 | 32.61 |
|  |  | R: TCCTCAGGCTTCAGTTTAAGCAT |  |  |  |  |  |  |  |  |  |
|  |  | P: TGGCCACTGTGTCTC |  |  |  |  |  |  |  |  |  |
|  | *esr2a* | F: TTTGTGGACCTGTGCCTGTTC | 118.3 | 25.09 | 23.36 | 25.10 | 25.98 | 24.68 | 24.79 | 26.46 | 26.06 |
|  |  | R: CATGAGCCCTAGCATCAGCA |  |  |  |  |  |  |  |  |  |
|  |  | P: TTGGAGTGCTGCTGGT |  |  |  |  |  |  |  |  |  |
|  | *esr2b* | F: AACGAGGCCTGTCATTCCAG | 102.9 | 22.77 | 19.57 | 22.11 | 19.99 | 21.58 | 22.19 | 22.66 | 20.31 |
|  |  | R: TGTGTGAGAGCAGCATGAGGA |  |  |  |  |  |  |  |  |  |
|  |  | P: TCATCCCGCCTGGC |  |  |  |  |  |  |  |  |  |
|  | *ar* | F: AAGTGGTCAAGTGGGCCAAA | 103.8 | 25.39 | 24.51 | 26.23 | 23.58 | 27.51 | 26.92 | 26.87 | 25.82 |
|  |  | R: ACCCCCATCCATGAATGCT |  |  |  |  |  |  |  |  |  |
|  |  | P: TTGCCAGGTTTTCGGAAT |  |  |  |  |  |  |  |  |  |
|  | *cyp19a1b* | F: CGGACGACGTGAGACAGTGT | 107.9 | 27.86 | 27.76 | - | 28.86 | 30.09 | 31.48 | - | - |
|  |  | R: TCCTCATCTCCACCTCAGGG |  |  |  |  |  |  |  |  |  |
|  |  | P: TGATAGCAGCCCCAGACA |  |  |  |  |  |  |  |  |  |
| Circadian rhythm | *nr1d1* | F: TGTCCCGCGATGCTGTT | 103.8 | 25.55 | 24.97 | 28.65 | 21.13 | 25.43 | 28.08 | 26.40 | 24.60 |
|  |  | R: CCAGCATGCGCTGCTTCT |  |  |  |  |  |  |  |  |  |
|  |  | P: CTTTGGCAGAATACC |  |  |  |  |  |  |  |  |  |
|  | *nr1d2a* | F: TCGGATAGCGGCGATGA | 112.3 | 23.22 | 23.92 | 25.27 | 21.46 | 26.79 | 27.07 | - | - |
|  |  | R: GAATGTGCCCTGGTGACACA |  |  |  |  |  |  |  |  |  |
|  |  | P: AGGTTCCTCCTTCCAGG |  |  |  |  |  |  |  |  |  |
|  | *nr1d2b* | F: TCGGCATGTCCAAGGACTCT | 99 | 26.69 | 25.40 | 28.16 | 25.27 | 25.34 | 28.21 | 29.16 | 26.65 |
|  |  | R: CGCTGCTTCTCACGCTTTG |  |  |  |  |  |  |  |  |  |
|  |  | P: TGCGTTTCGGCCGCA |  |  |  |  |  |  |  |  |  |
|  | *arntl1a* | F: TCCAGCAGCCCGAGTAATG | 104.1 | 27.76 | 26.99 | 29.59 | 26.29 | 25.54 | 28.54 | 28.39 | 28.19 |
|  |  | R: GCCTCCAGAAGGCTCATGATC |  |  |  |  |  |  |  |  |  |
|  |  | P: CGAGGCAGCCATGG |  |  |  |  |  |  |  |  |  |
|  | *per1a/b* | F: CTGGACAACATTGCCTCAGAGTAC | 107 | 29.79 | 28.03 | 27.22 | 27.76 | 27.54 | 27.67 | - | - |
|  |  | R: GAAGGACACCGCCATGGA |  |  |  |  |  |  |  |  |  |
|  |  | P: CCCTCAAAAATACAGACACC |  |  |  |  |  |  |  |  |  |
|  | *per2* | F: GAGTACACCCTCAAAAACAATGACA | 106.5 | 28.37 | 29.05 | - | 27.50 | 29.10 | 30.42 | 28.03 | 25.90 |
|  |  | R: ATGTAGACGATCTTCCCCGTGAT |  |  |  |  |  |  |  |  |  |
|  |  | P: TGCGGTAGCGATATC |  |  |  |  |  |  |  |  |  |
|  | *cry1a/b* | F: CCGCCGGGACAAGGA | 99.1 | 25.40 | 26.33 | 28.91 | 24.44 | 25.52 | 28.60 | 26.93 | 24.40 |
|  |  | R: AGATGATTTCTACTCCGTGCTCTTC |  |  |  |  |  |  |  |  |  |
|  |  | P: TGGGCCGGTTAGC |  |  |  |  |  |  |  |  |  |
|  | *cry2* | F: TCCGCCTCTTCCTGAATGG | 107.5 | 27.01 | 25.40 | 29.15 | 25.02 | 27.47 | 29.12 | 27.85 | 27.18 |
|  |  | R: CCCACTACAGAGGCCTCAGTCT |  |  |  |  |  |  |  |  |  |
|  |  | P: TGGCATCTGTGCCC |  |  |  |  |  |  |  |  |  |
|  | *roraa/b* | F: TGCCGTGCCTTCGACTCT | 109.2 | 26.26 | 26.28 | 29.03 | 28.73 | 26.75 | 28.34 | 29.93 | 26.85 |
|  |  | R: CAGGCCCAGCATACTTTCCA |  |  |  |  |  |  |  |  |  |
|  |  | P: ACAACACAGTCTATTTCG |  |  |  |  |  |  |  |  |  |
|  | *clocka* | F: CGCAATGCAGCACCTGAA | 104.5 | 25.15 | 25.23 | 26.73 | 26.31 | 24.73 | 26.91 | 27.31 | 25.79 |
|  |  | R: ATGTTGGCCTCGATCATCCT |  |  |  |  |  |  |  |  |  |
|  |  | P: CTGGAGCAGAGGAC |  |  |  |  |  |  |  |  |  |
|  | *clockb* | F: CCTGGAGTCACTGGCCAAAT | 106.9 | 25.86 | 25.61 | 28.51 | 26.37 | 26.26 | 30.02 | 30.30 | 28.46 |
|  |  | R: CACGACTTCCCCTTCCCATA |  |  |  |  |  |  |  |  |  |
|  |  | P: CCACGAACACTTAATG |  |  |  |  |  |  |  |  |  |
|  | *npas2* | F: ACGCATGCTGCTCAACCA | 110.5 | 26.82 | 26.70 | 28.90 | 25.48 | 25.75 | 32.02 | - | - |
|  |  | R: CCTGGGAGCTAACACGGCTAT |  |  |  |  |  |  |  |  |  |
|  |  | P: CAGTGCAGACACTCGTA |  |  |  |  |  |  |  |  |  |
|  | *dbpa/b* | F: GAAGGGAGACCGTGCAAGTC | 107.5 | 25.49 | 27.23 | 27.59 | 25.94 | 31.28 | 31.66 | 35.22 | 32.47 |
|  |  | R: GGAGTCCTCATCCATGTCTGTGA |  |  |  |  |  |  |  |  |  |
|  |  | P: TGCGACGTTAAAGAC |  |  |  |  |  |  |  |  |  |
| Apoptosis | *casp3a/b* | F: AAAGGGATAGCTGCACAAGGG | 107.7 | 23.51 | 23.67 | 29.52 | 25.14 | 23.03 | 26.57 | - | - |
|  |  | R: TGTGTACATGTCAGCCGGAGG |  |  |  |  |  |  |  |  |  |
|  |  | P: CAGTCCCTCAGTGCAAA |  |  |  |  |  |  |  |  |  |
|  | *casp9* | F: GCCAGACAGTTGGTTCGAGAC | 94.6 | 30.41 | 28.90 | 31.27 | 29.21 | 27.19 | 30.46 | 33.00 | 28.92 |
|  |  | R: GGCTATGCTGCCCTTTCTCA |  |  |  |  |  |  |  |  |  |
|  |  | P: TCCCAGCTTTAATAGAG |  |  |  |  |  |  |  |  |  |
|  | *tp53* | F: CCTTTGAGGTGCGTGTGTGT | 108.8 | - | - | 26.61 | 25.17 | 24.61 | 27.32 | 27.07 | 25.55 |
|  |  | R: GGGTTGTCTCCTGCTGCTTCT |  |  |  |  |  |  |  |  |  |
|  |  | P: CTGGTCGAGACAGGAA |  |  |  |  |  |  |  |  |  |
|  | *pdcd10a/b* | F: AGGCTGAGAAGGAGAACCCAG | 106.5 | 25.83 | 24.84 | 28.15 | 24.83 | 23.95 | 27.68 | 29.64 | 26.24 |
|  |  | R: CACATCGTCTGCAGCCATTC |  |  |  |  |  |  |  |  |  |
|  |  | P: TGACCCAGGACATCAT |  |  |  |  |  |  |  |  |  |
|  | *chmp5a/b* | F: CCAAAGAGATGAAGGCGGC | 127.8 | 23.77 | 23.25 | 25.42 | 21.53 | 22.45 | 25.72 | 26.61 | 23.31 |
|  |  | R: TTGGCGTCCTCCATCATGT |  |  |  |  |  |  |  |  |  |
|  |  | P: AGGATCTCCAGGACCAG |  |  |  |  |  |  |  |  |  |
| Growth and metabolism | *ghr* | F: TAACCGGGAGCCACTTTGAC | 105.5 | 23.81 | 24.34 | 24.59 | 23.03 | 25.28 | 26.46 | 26.24 | 24.23 |
|  |  | R: ATTGACCTCACGGTACTGCACC |  |  |  |  |  |  |  |  |  |
|  |  | P: TGAGCTGGGAGCCG |  |  |  |  |  |  |  |  |  |
|  | *igf1* | F: TTCAAGAGTGCGATGTGCTGT | 108.4 | 22.59 | 20.41 | 22.70 | 22.72 | 24.02 | 21.83 | 25.53 | 25.83 |
|  |  | R: CGCCGAAGTCAGGGTTAGG |  |  |  |  |  |  |  |  |  |
|  |  | P: ACACCCTCTCACTGCT |  |  |  |  |  |  |  |  |  |
|  | *igf2* | F: ATGTGGAGGAGAACTGGTGGAC | 94.1 | 25.05 | 23.36 | 25.60 | 24.03 | 25.41 | 24.94 | 25.70 | 25.11 |
|  |  | R: CCTGCTGGTTGGCCTACTGA |  |  |  |  |  |  |  |  |  |
|  |  | P: TGCAGTTCGTCTGTGAAG |  |  |  |  |  |  |  |  |  |
|  | *igfbp1* | F: AACTGTGCGGAATCTACACGG | 99.2 | 29.61 | 29.71 | 28.96 | 24.19 | 29.36 | - | - | - |
|  |  | R: CTGGCTGCGAATAAGGGAGT |  |  |  |  |  |  |  |  |  |
|  |  | P: TGCACCCCAATACC |  |  |  |  |  |  |  |  |  |
|  | *igfbp2* | F: AGCTGCATGTCCTAAGCTTGC | 99.3 | 24.55 | 23.83 | 25.47 | 23.12 | 25.35 | 31.61 | 25.53 | 25.25 |
|  |  | R: GCCTTCTAACCGGGAACACA |  |  |  |  |  |  |  |  |  |
|  |  | P: TCAGGGAGCCCGGCT |  |  |  |  |  |  |  |  |  |
|  | *ampka1* | F: AAGTTTGAGTGCACCGAGGAG | 105.3 | 25.81 | 24.07 | 27.03 | 24.31 | 24.40 | 25.77 | 25.83 | 24.01 |
|  |  | R: GACATAATGCGGCGGTTGTC |  |  |  |  |  |  |  |  |  |
|  |  | P: CGCAACCACCACGAC |  |  |  |  |  |  |  |  |  |
|  | *ldhb* | F: CCCCCAACTGCACCCTTATT | 96.7 | - | - | 24.64 | 24.76 | 24.68 | 26.49 | 24.07 | 23.31 |
|  |  | R: AATCCGCTCAACTTCCACGT |  |  |  |  |  |  |  |  |  |
|  |  | P: CCAACCCAGTGGACGT |  |  |  |  |  |  |  |  |  |
|  | *pck1* | F: ATATGAGAACTGCTGGCTGGC | 102.1 | 30.24 | - | 30.87 | 23.33 | 29.60 | - | 30.89 | 27.64 |
|  |  | R: CACCTACTCGTGGAGACGGAA |  |  |  |  |  |  |  |  |  |
|  |  | P: CCCAGAGACGTGGCC |  |  |  |  |  |  |  |  |  |
|  | *fasn* | F: TGTGGGAGGTGTAGTCAAGCC | 114.9 | 23.17 | 19.02 | 23.97 | 24.85 | 19.98 | 18.42 | 20.45 | 18.33 |
|  |  | R: TCCCTGGGCCATGTATCTGA |  |  |  |  |  |  |  |  |  |
|  |  | P: AGGTGGAGGAGGCC |  |  |  |  |  |  |  |  |  |
|  | *cpt1a* | F: TACAGCTGGCCCAATTCAGG | 112 | 24.37 | 26.52 | 27.73 | 24.02 | 27.24 | 29.87 | 28.99 | 23.63 |
|  |  | R: AACCGTCTCTGTCCTACCCTCA |  |  |  |  |  |  |  |  |  |
|  |  | P: TCAATGACCCGGATGTT |  |  |  |  |  |  |  |  |  |
|  | *cpt1b* | F: GGACCAGTCCTGATGCCTTC | 110 | 26.05 | 27.46 | 28.25 | 24.71 | 26.76 | 29.75 | - | 23.71 |
|  |  | R: TCGAGGCCTCATACGTCAGAC |  |  |  |  |  |  |  |  |  |
|  |  | P: ACTGCAGCTGGCTCA |  |  |  |  |  |  |  |  |  |
|  | *cs* | F: TTGATTGCCAAGTTGCCGT | 116 | 24.30 | 23.79 | 25.01 | 22.80 | 21.90 | 24.43 | 23.12 | 22.86 |
|  |  | R: ATGTTGGCGAAGTTAGCGGA |  |  |  |  |  |  |  |  |  |
|  |  | P: CAGCAGCATTGGC |  |  |  |  |  |  |  |  |  |
|  | *ldha* | F: CGTCAAGTACAGCCCCAACG | 104 | - | 28.41 | 30.74 | - | 30.85 | 27.26 | 27.44 | 31.04 |
|  |  | R: GCCACGTAGGTCAGGATGTCA |  |  |  |  |  |  |  |  |  |
|  |  | P: TGCTGGTCGTCTCCAA |  |  |  |  |  |  |  |  |  |
|  | *lipea/b* | F: ACACTGGTCAAGGTGTTGCAGT | 121 | 28.06 | 27.57 | 27.83 | 26.63 | 26.40 | 27.62 | 30.15 | 27.13 |
|  |  | R: TGCAGTTAGCGGCAATGTAGC |  |  |  |  |  |  |  |  |  |
|  |  | P: CTTCTGCACATCATCCA |  |  |  |  |  |  |  |  |  |
|  | *lpl* | F: TGACAGCGCTGTACAAGAGGG | 111 | 22.82 | 22.75 | 25.32 | 26.01 | 23.35 | 24.90 | - | 25.83 |
|  |  | R: GGAGGTGAGGTAGTGCTGCTG |  |  |  |  |  |  |  |  |  |
|  |  | P: TGGACTGGCTGACACGG |  |  |  |  |  |  |  |  |  |
|  | *ctsd* | F: TTCACAGACATCGCCTGCTT | 126 | 23.95 | 22.61 | 23.05 | 22.88 | 21.52 | 22.49 | 23.72 | 23.46 |
|  |  | R: GGTACCCAGACAGACTGCCAG |  |  |  |  |  |  |  |  |  |
|  |  | P: CCACAAGTATAACGGTGCC |  |  |  |  |  |  |  |  |  |
| Detoxification | *cyp1a* | F: TACAGCTGGCCCAATTCAGG | 98.9 | 25.29 | 22.34 | 23.32 | 25.27 | 22.97 | 23.98 | 24.36 | 25.58 |
|  |  | R: AACCGTCTCTGTCCTACCCTCA |  |  |  |  |  |  |  |  |  |
|  |  | P: TCAATGACCCGGATGTT |  |  |  |  |  |  |  |  |  |
|  | *gstp1* | F: GGTGACAAGCCTTCGTTTGC | 110 | 23.95 | 20.61 | 23.56 | 24.02 | 21.04 | 22.74 | 24.58 | 23.26 |
|  |  | R: CAAAGCTCTTCAGGGAGGGG |  |  |  |  |  |  |  |  |  |
|  |  | P: TGAAGTGCTGCTCAAC |  |  |  |  |  |  |  |  |  |
|  | *gpx1a* | F: CCACCCCTTGTTTGTGTATCTCA | 119.1 | - | 24.10 | 27.34 | 24.77 | 23.12 | 26.39 | - | - |
|  |  | R: GGGCTCCACATGATGAACTTG |  |  |  |  |  |  |  |  |  |
|  |  | P: CCATTCCCCTCCGATG |  |  |  |  |  |  |  |  |  |
|  | *cat* | F: TGGGCCGCTACAACAGTACTG | 103.8 | 21.35 | 20.87 | 23.92 | 23.12 | 22.73 | 24.00 | 23.18 | 21.52 |
|  |  | R: CTCGTTCAGCACCTTAGTGAAGAA |  |  |  |  |  |  |  |  |  |
|  |  | P: CGTCACACAGGTGCGTA |  |  |  |  |  |  |  |  |  |
|  | *glul* | F: TGAAGTCATGCCTGCACAGTG | 95.1 | 23.07 | 23.04 | 25.77 | 22.24 | 24.45 | 26.09 | 25.94 | 24.51 |
|  |  | R: CACACCCGGTGGAGAATGA |  |  |  |  |  |  |  |  |  |
|  |  | P: CAGGTTGGCCCTTGT |  |  |  |  |  |  |  |  |  |
|  | *sod2* | F: GATGGCTGGGCTTTGACAA | 105.3 | 23.29 | 22.48 | 25.61 | 23.23 | 22.97 | 25.35 | 23.86 | 21.35 |
|  |  | R: CCTGCAGTGGGTCTTGATTAGG |  |  |  |  |  |  |  |  |  |
|  |  | P: AAGCTCCGTATCACAGC |  |  |  |  |  |  |  |  |  |
|  | *sod1* | F: CAACACCAACGGCTGTATGAGT | 102.2 | 29.65 | 20.85 | 22.42 | 22.97 | 21.46 | 22.42 | 22.14 | 20.13 |
|  |  | R: CCTCCGTGGGTCTTGTTGTG |  |  |  |  |  |  |  |  |  |
|  |  | P: CGGACCCCACTTCA |  |  |  |  |  |  |  |  |  |
|  | *nfe2l2a* | F: GGTGGCTACAGCGATTCAGAC | 99.9 | 25.68 | 24.58 | 27.76 | 25.76 | 25.00 | 26.13 | 26.50 | - |
|  |  | R: GAAAGAGGTGTCTGCAGGCC |  |  |  |  |  |  |  |  |  |
|  |  | P: AGTAACCCTGGGAGTGC |  |  |  |  |  |  |  |  |  |
| Hypoxia | *hif1a* | F: GCTGTGGGCTGAAGAGTGATC | 122 | 27.91 | 28.26 | 27.57 | 26.76 | 28.06 | 26.85 | 27.46 | 26.47 |
|  |  | R: CTGGGTTCTCCTTCAGCTGG |  |  |  |  |  |  |  |  |  |
|  |  | P: TCTCTGAGGCCTCCGAG |  |  |  |  |  |  |  |  |  |
|  | *epor* | F: TAAAGTGGCTCTGCTGCTCCA | 111.6 | 27.53 | 28.20 | 28.91 | 27.44 | 28.87 | 30.50 | 34.14 | - |
|  |  | R: CCAGAAGCAGGTGAGGTCTGA |  |  |  |  |  |  |  |  |  |
|  |  | P: CTGAGCCAGAGAATC |  |  |  |  |  |  |  |  |  |
|  | *vegfc* | F: AACCACACCGTGTGCAATTG | 103.3 | 28.72 | 28.18 | 27.47 | 26.52 | 26.90 | 29.40 | - | - |
|  |  | R: TGTGGTTCTTTGGGCAGGTT |  |  |  |  |  |  |  |  |  |
|  |  | P: ACTGGACGCCTACAGAC |  |  |  |  |  |  |  |  |  |
|  | *slc2a1a* | F: TGAGCATCGTGGCCATCTT | 121.3 | - | 24.26 | 27.48 | 26.24 | 25.20 | 25.47 | 23.66 | - |
|  |  | R: AACAGCTCAGCCACGATGAA |  |  |  |  |  |  |  |  |  |
|  |  | P: TTGGCCCGGGTCC |  |  |  |  |  |  |  |  |  |
|  | *mb* | F: ATGGTTCTGAAGTGCTGGGG | 102.6 | - | - | - | - | 29.57 | 30.58 | 27.09 | 28.29 |
|  |  | R: AAACAGACGGCTCAGAACCAG |  |  |  |  |  |  |  |  |  |
|  |  | P: AGGCTGACTACAACAAA |  |  |  |  |  |  |  |  |  |
|  | *hk1* | F: ATGCTGGAGGACATTCGCA | 124.7 | 28.90 | 27.89 | - | - | - | 28.60 | 28.06 | 28.88 |
|  |  | R: TTCGCCCATGTACATCCCA |  |  |  |  |  |  |  |  |  |
|  |  | P: AGGGGCTCTCTAAAC |  |  |  |  |  |  |  |  |  |
|  | *pfkm* | F: TGGTATCTACACCGGAGCCAA | 126 | - | - | 28.50 | - | 30.87 | 27.85 | 30.06 | - |
|  |  | R: TGCAGCATCATGGACACACTC |  |  |  |  |  |  |  |  |  |
|  |  | P: CCAGGGTCTGGTGGAT |  |  |  |  |  |  |  |  |  |
|  | *aldoaa* | F: CAACGGAGAGACCACCACTCA | 102.1 | 26.12 | 25.43 | 26.51 | 25.53 | 25.16 | 26.82 | 24.79 | 26.48 |
|  |  | R: ACGCCACTTAGCAAAGTCAGC |  |  |  |  |  |  |  |  |  |
|  |  | P: TGTACGAGCGGTGTGC |  |  |  |  |  |  |  |  |  |
|  | *eno1a* | F: TCCCTGCCTTCAACGTGATC | 112.2 | 22.05 | 21.25 | 22.39 | 22.01 | 21.56 | 21.55 | 22.32 | 21.31 |
|  |  | R: CCTCATGGCCTCCTTGAAGG |  |  |  |  |  |  |  |  |  |
|  |  | P: CGGAGGTTCCCACGCA |  |  |  |  |  |  |  |  |  |
|  | *pgk1* | F: AGATGATCATCGGTGGTGGC | 122.4 | 23.77 | 22.79 | 24.95 | 22.98 | 23.40 | 24.97 | 24.62 | 23.53 |
|  |  | R: CATACAGGGAGGTGCCGATC |  |  |  |  |  |  |  |  |  |
|  |  | P: CCTTCACCTTCCTCAATG |  |  |  |  |  |  |  |  |  |
| Endogenous control | *rpl7* | F: TCGTCATCAGGATCAGGGGT | 116.2 | 17.78 | 17.90 | 18.94 | 18.26 | 17.70 | 19.11 | 19.78 | 19.46 |
|  |  | R: GAAGATCTGACGCAGACGCA |  |  |  |  |  |  |  |  |  |
|  |  | P: CAAGGTGCGCAAGGT |  |  |  |  |  |  |  |  |  |
|  | *rps9* | F: TTCTCCCTGCGTTCACCATAC | 109.9 | 19.97 | 19.63 | 22.05 | 18.39 | 19.61 | 20.72 | 20.70 | 19.15 |
|  |  | R: GGCCCTTCTTGGCATTCTTT |  |  |  |  |  |  |  |  |  |
|  |  | P: CCCGGCCGTGTCAA |  |  |  |  |  |  |  |  |  |
|  | *ef1a* | F: GAAGCTTGAGGACAACCCCA | 99.9 | 23.05 | 23.70 | 25.23 | 24.19 | 23.36 | 22.80 | 23.75 | 24.39 |
|  |  | R: GAAGCTCTCCACACACATGGG |  |  |  |  |  |  |  |  |  |
|  |  | P: CCGCCATCATCGTCAT |  |  |  |  |  |  |  |  |  |
|  | *rpl13a* | F: CACTGGAGAGGCTGAAGGTGT | 108.5 | 19.23 | 19.81 | 20.43 | 18.43 | 19.63 | 21.41 | 20.77 | 20.57 |
|  |  | R: GTGGGCTTCAGACGGACAAT |  |  |  |  |  |  |  |  |  |
|  |  | P: CATGGTCGTACCTGCT |  |  |  |  |  |  |  |  |  |

Supplementary Table 3: TaqMan primer and probe sequences, PCR efficiencies, and mean C_T_ levels for the 112 candidate genes for muscle tissue across eight salmonid species

| Biological function | Gene symbol | Primer and probe sequences | Efficiency (%) | C_T_ level | | | | | | | |
| --- | --- | --- | --- | --- | --- | --- | --- | --- | --- | --- | --- |
|  |  |  |  | *S. salar* | *S. trutta* | *O. mykiss* | *O. tshawytscha* | *S. fontinalis* | *S. alpinus* | *C. clupeaformis* | *C. hoyi* |
| Stress | *gr1* | F: GTCTTTGGCCTGTATCCCCC | 88.8 | 26.47 | 24.78 | 26.34 | 24.46 | - | - | 26.60 | 26.02 |
|  |  | R: AGCTCGACATCCCTGATCCA |  |  |  |  |  |  |  |  |  |
|  |  | P: TGCCCTCGGTCAGTGA |  |  |  |  |  |  |  |  |  |
|  | *gr2* | F: ATGGCAGACCAGTGTGAACAGAT | 107.6 | 29.19 | 26.60 | 28.25 | 25.01 | 27.22 | 28.62 | 26.51 | 27.05 |
|  |  | R: GAGCAGCAGCAGAACCTTCAT |  |  |  |  |  |  |  |  |  |
|  |  | P: AGACTGCAGGTGTCTCA |  |  |  |  |  |  |  |  |  |
|  | *mr* | F: CATAGTCAATGTCAGCTGCTCCC | 94.9 | 26.97 | 24.41 | 25.91 | 24.04 | 26.10 | 26.57 | 26.79 | 25.88 |
|  |  | R: GCTGCTGCTCTGGCTTCTTCT |  |  |  |  |  |  |  |  |  |
|  |  | P: CCAGCAGCAACACA |  |  |  |  |  |  |  |  |  |
|  | *hsd11b2* | F: CTGTCTAGCAGCGTACGGAGC | 100.4 | 28.97 | 26.10 | 27.21 | 24.67 | 28.03 | 27.34 | 26.62 | 27.45 |
|  |  | R: ATGGTGGACACTTTGACCCC |  |  |  |  |  |  |  |  |  |
|  |  | P: CCTGTTCATCAACACACT |  |  |  |  |  |  |  |  |  |
|  | *hsp70a* | F: GAGAACACTGTCCTCCAGCTCC | 100 | 25.54 | 26.63 | 27.71 | 24.83 | 26.63 | 29.24 | 26.22 | 27.34 |
|  |  | R: CCCTGAAGAGGTCGGAACAC |  |  |  |  |  |  |  |  |  |
|  |  | P: ACACCTCCATCACCAGG |  |  |  |  |  |  |  |  |  |
|  | *hsp90aa* | F: AAGATCGAGGTCACCCCTGA | 102.5 | 23.40 | 23.20 | 24.43 | 22.67 | 22.59 | 27.20 | 32.33 | 31.46 |
|  |  | R: GGTGCCAGACTTTGCAATGG |  |  |  |  |  |  |  |  |  |
|  |  | P: CGGCATCGGCATGA |  |  |  |  |  |  |  |  |  |
|  | *hsp90ba* | F: GAGGTGGAGGAGGACGAGTACA | 100.4 | 29.89 | 26.67 | 29.28 | 27.36 | 26.73 | 27.67 | 28.49 | 26.84 |
|  |  | R: GCTGTGAAGTGGATGTGGGA |  |  |  |  |  |  |  |  |  |
|  |  | P: TCTCCAGGGACACAGAC |  |  |  |  |  |  |  |  |  |
|  | *hsf1* | F: CCCAAGTTCAGCAGGCAGTAC | 93.8 | 25.38 | 23.08 | 24.99 | 24.60 | 24.32 | 25.56 | 24.61 | 25.34 |
|  |  | R: GCCGTGAAGAGACCGGTACT |  |  |  |  |  |  |  |  |  |
|  |  | P: TGCAGGGCTCGCCT |  |  |  |  |  |  |  |  |  |
|  | *cirbpa* | F: CGGGAAGGTCTCGTGGATT | 109.3 | 24.07 | 20.24 | 25.45 | 22.63 | 22.48 | 24.54 | 22.46 | 22.76 |
|  |  | R: TGGTTCTGCCATCGACAGACT |  |  |  |  |  |  |  |  |  |
|  |  | P: CATTGGAGGGAATGAA |  |  |  |  |  |  |  |  |  |
|  | *crfb1* | F: TCATTGCTTTCTTACCGCGC | 100.5 | 28.85 | - | - | 28.20 | 28.94 | - | - | - |
|  |  | R: AGGAGGGGAAGAGACTGTTGCT |  |  |  |  |  |  |  |  |  |
|  |  | P: CCGCTCCACATCACGA |  |  |  |  |  |  |  |  |  |
|  | *avt* | F: CTAGACCCAGACTGCCTAGAGGAC | 96.9 | 30.73 | 29.88 | 32.64 | 28.99 | 30.03 | - | - | - |
|  |  | R: GCCAAACCACCCATTAAGGC |  |  |  |  |  |  |  |  |  |
|  |  | P: ACGTCAGTCACCCAGCGA |  |  |  |  |  |  |  |  |  |
|  | *mmp2* | F: CGCTGTGGAGTTCCTGATGTT | 99.6 | 25.81 | 22.09 | 24.23 | 20.83 | 24.60 | 25.17 | 25.31 | 23.86 |
|  |  | R: AGGTCAGGAGAGTGGCCTAGAA |  |  |  |  |  |  |  |  |  |
|  |  | P: AGGAAACCCAAGTGGCA |  |  |  |  |  |  |  |  |  |
|  | *mmp9* | F: ACGGCAAAGCATGTGTGTTC | 105.1 | 29.58 | 26.42 | 28.75 | 28.08 | 28.68 | 29.31 | - | - |
|  |  | R: GGAACACCAGCGGTATCCAT |  |  |  |  |  |  |  |  |  |
|  |  | P: AGGTTGCACGACGGAA |  |  |  |  |  |  |  |  |  |
|  | *mt* | F: GCTCAAAAACTGGACGCTGC | 116.8 | 26.82 | 29.30 | - | 28.93 | - | - | - | - |
|  |  | R: GGCAGCAGGAACAACAACTTT |  |  |  |  |  |  |  |  |  |
|  |  | P: TACAAACTGCGGATGTGC |  |  |  |  |  |  |  |  |  |
|  | *mtA* | F: TGGATCCTTGTGAATGCTCCA | 105.1 | 26.80 | 22.41 | 26.31 | 23.21 | 25.38 | 26.85 | 25.27 | 24.74 |
|  |  | R: CTTACAACTGGTGCATGCGC |  |  |  |  |  |  |  |  |  |
|  |  | P: CGGTGGATCCTGCAAG |  |  |  |  |  |  |  |  |  |
|  | *mtB* | F: GAAAAGTTGCTGCCCCTGC | 99.9 | 27.79 | 27.79 | 26.20 | 23.56 | 27.99 | 26.75 | 27.97 | 26.01 |
|  |  | R: AACAGCTGGTATCGCAGGTCTT |  |  |  |  |  |  |  |  |  |
|  |  | P: TTCAGGCTGTGTGTGCA |  |  |  |  |  |  |  |  |  |
|  | *hsd20b2* | F: ACATTGTACTGGTCAGCCGGT | 107.4 | 28.45 | 29.77 | - | 30.13 | 28.72 | 28.90 | 29.41 | 29.92 |
|  |  | R: TGGCCCTCTGTGAAGTCTGTC |  |  |  |  |  |  |  |  |  |
|  |  | P: CCAACATGGACGCCAGA |  |  |  |  |  |  |  |  |  |
|  | *hsp30* | F: TGCTGTGTTCCCGAGGATTC | 113.1 | 23.20 | 23.15 | 29.50 | 28.24 | 27.09 | 28.52 | 28.18 | 23.00 |
|  |  | R: TCTCTGCAGTAGATCCCGCTG |  |  |  |  |  |  |  |  |  |
|  |  | P: TGTGCGCAGTCTAT |  |  |  |  |  |  |  |  |  |
|  | *serpinh1a* | F: AGCGCTGTGAAGTCCATCAA | 96.5 | 26.33 | 23.04 | 25.99 | 24.02 | 23.79 | 25.13 | 30.13 | 27.46 |
|  |  | R: TGATGATCATGGCCCCATC |  |  |  |  |  |  |  |  |  |
|  |  | P: CGGCCAAGTCCACCGA |  |  |  |  |  |  |  |  |  |
|  | *hspa4* | F: ACTGCTGAGACCGCAATGAA | 112 | 25.30 | 23.86 | 27.19 | 24.19 | 24.95 | 26.16 | 25.26 | 25.17 |
|  |  | R: TGCGTCAGTGTAGAAGCTGGG |  |  |  |  |  |  |  |  |  |
|  |  | P: ACCTGTGGCTGATTGT |  |  |  |  |  |  |  |  |  |
|  | *hsp7c* | F: TAGCCAACGACCAGGGAAAC | 95.1 | 24.10 | 23.38 | 28.12 | 23.80 | 24.02 | 28.19 | 28.19 | 30.13 |
|  |  | R: CGTCCCCAATCAGCCTTTCT |  |  |  |  |  |  |  |  |  |
|  |  | P: ACCCAGTTATGTCGCCTT |  |  |  |  |  |  |  |  |  |
|  | *pomc-a1* | F: AGAACAGCATCCTGGAGTGCA | 124.7 | - | - | - | 30.22 | - | - | - | - |
|  |  | R: GGGAGTTGGGTTGGAGATGG |  |  |  |  |  |  |  |  |  |
|  |  | P: ACCTCACCGCCGAAT |  |  |  |  |  |  |  |  |  |
| Neural plasticity | *bdnf* | F: TCCTTCCTCTGTGGTCACCAC | 88.8 | 30.57 | 30.42 | - | 31.32 | 31.66 | - | - | - |
|  |  | R: AGGCACTTGGTTGCTGATCA |  |  |  |  |  |  |  |  |  |
|  |  | P: TGTCGACCTGTATGCAT |  |  |  |  |  |  |  |  |  |
|  | *pcna* | F: TGGAGGCTCTGAAGGACCTG | 109 | 28.30 | 24.76 | 27.59 | 26.28 | 25.97 | 27.51 | 28.99 | 26.51 |
|  |  | R: GAGCTGAACTAGGGAGACGTGG |  |  |  |  |  |  |  |  |  |
|  |  | P: CTGTTGGGACGTGAGCT |  |  |  |  |  |  |  |  |  |
|  | *neurod1* | F: ATGTCATCAGACCCCCAGGA | 112.3 | - | 30.92 | - | 29.63 | - | - | - | - |
|  |  | R: GATTCGAGTGCGCCGTTTT |  |  |  |  |  |  |  |  |  |
|  |  | P: CGACCTGGACTGATGA |  |  |  |  |  |  |  |  |  |
|  | *dcx* | F: GTCTGTGAACGTGAAGGCCTC | 99 | 31.58 | 26.88 | 31.00 | 27.49 | 29.11 | 29.53 | 30.67 | 28.55 |
|  |  | R: TTGGGCCGAACAAACTCCT |  |  |  |  |  |  |  |  |  |
|  |  | P: AGCCAGAAGAACCTG |  |  |  |  |  |  |  |  |  |
|  | *neurog1* | F: AGAGCGAGGAATGAAGCCAC | 77.1 | - | 24.83 | - | 30.89 | 24.98 | - | - | - |
|  |  | R: GTTTCTCTCGCGGTCGTTTG |  |  |  |  |  |  |  |  |  |
|  |  | P: CGTGGTCAAGAAGAA |  |  |  |  |  |  |  |  |  |
|  | *sox2* | F: TCAGACAGGAGACCTACGGGAC | 104.9 | - | 28.57 | - | 27.85 | 29.10 | - | 30.68 | 29.82 |
|  |  | R: TTGGGACATGTGGAGTCTGC |  |  |  |  |  |  |  |  |  |
|  |  | P: TTACCGGGCGCTGAG |  |  |  |  |  |  |  |  |  |
| Appetite regulation | *agrp* | F: ATACTGCTGGACCCTGTGCCT | 126.8 | - | 31.51 | - | - | - | - | - | - |
|  |  | R: CGGGATGGGAGTCGTCTAGA |  |  |  |  |  |  |  |  |  |
|  |  | P: CCTGATCCAGCTGGCTA |  |  |  |  |  |  |  |  |  |
|  | *cart* | F: TGACCAACGAAAAGCAACTGC | 93.9 | - | 31.50 | - | - | - | - | - | - |
|  |  | R: AGGGACTTGGCCGAATTTCT |  |  |  |  |  |  |  |  |  |
|  |  | P: CTGCAGACCAAGAGA |  |  |  |  |  |  |  |  |  |
|  | *cck-I* | F: GGGTCCCAGCCACAAGATAA | 97.7 | 30.48 | 28.12 | - | 27.53 | - | 29.25 | - | - |
|  |  | R: CTCGTACTCCTCTGCACTGCG |  |  |  |  |  |  |  |  |  |
|  |  | P: TGGGCTGGATGGAC |  |  |  |  |  |  |  |  |  |
|  | *lepr* | F: CGCTGTATGCACATCAACGG | 94.6 | 30.09 | 28.64 | 29.82 | 26.57 | 28.66 | 30.25 | 26.39 | 29.31 |
|  |  | R: GTCAGGAGCTCTGCTGTTGTGA |  |  |  |  |  |  |  |  |  |
|  |  | P: CCGGGACCTGGAGTGA |  |  |  |  |  |  |  |  |  |
|  | *lepa* | F: AAGCCTGCCTTCCATAGTGGA | 119.6 | - | 30.95 | - | 33.23 | - | - | 30.93 | 33.15 |
|  |  | R: TGAGGTCTGCCCAGTCTAGGA |  |  |  |  |  |  |  |  |  |
|  |  | P: TGGGATTCTACCAGGACC |  |  |  |  |  |  |  |  |  |
|  | *npy* | F: AACCTCATCACAAGGCAGAGGT | 104.6 | 30.56 | 28.57 | - | 29.56 | - | - | - | - |
|  |  | R: ACGTGTCTGTGCTCTCCTTCAG |  |  |  |  |  |  |  |  |  |
|  |  | P: AGGTCCAGCCCTGAC |  |  |  |  |  |  |  |  |  |
| Osmoregulation | *cftr* | F: CAGGCAGAAAAGAGCAGGGA | 127.9 | - | 27.10 | - | - | - | - | - | - |
|  |  | R: TCTCCATCACCTCCTCCCAG |  |  |  |  |  |  |  |  |  |
|  |  | P: TCGTCGCCTGGCTC |  |  |  |  |  |  |  |  |  |
|  | *atpa1a* | F: GGTGGTTGGAGATCTGGTGG | 115.3 | - | 28.31 | - | 26.13 | 26.04 | - | - | 27.92 |
|  |  | R: CACCAGTGAGGGAGGAGTTGTC |  |  |  |  |  |  |  |  |  |
|  |  | P: ATTTGCGTATTGTCTCTG |  |  |  |  |  |  |  |  |  |
|  | *atpa1b* | F: GGATGCTTGGCTGGGATTT | 98.3 | 25.09 | 22.87 | 27.64 | 24.63 | 24.20 | 25.73 | 26.09 | 24.77 |
|  |  | R: CTCTGTCCTGGTAGGTGGGTTT |  |  |  |  |  |  |  |  |  |
|  |  | P: CAGGCCCTGCTGCT |  |  |  |  |  |  |  |  |  |
|  | *crhbp* | F: GAGCCCAACCAGGTCATCAA | 81 | - | 29.53 | - | 28.13 | 29.40 | - | - | 31.24 |
|  |  | R: CTCTCCCTTCATCACCCAGC |  |  |  |  |  |  |  |  |  |
|  |  | P: CGTTGACATCGACTGCA |  |  |  |  |  |  |  |  |  |
| Immune function | *cam* | F: GAGGAGCAGATTGCCGAGTT | 105 | 24.55 | 20.72 | 24.85 | 21.49 | 22.03 | 23.44 | 23.24 | 21.49 |
|  |  | R: CATGACAGTGCCCAGCTCTTT |  |  |  |  |  |  |  |  |  |
|  |  | P: CGCTCTTTGACAAGGA |  |  |  |  |  |  |  |  |  |
|  | *mhc-I* | F: AGTCCCTCCCTCAGTGTCTCTG | 98.6 | 25.49 | 21.60 | 25.66 | 21.69 | 24.75 | 25.97 | 22.91 | 23.35 |
|  |  | R: AGGACACCATGACTCCACTGG |  |  |  |  |  |  |  |  |  |
|  |  | P: AGTGACCTGCCACGCG |  |  |  |  |  |  |  |  |  |
|  | *mhc-II* | F: CTCACAGCAGCATCTACCCCA | 88.4 | 34.78 | 31.38 | 28.61 | 26.81 | 28.81 | - | 27.18 | 24.60 |
|  |  | R: CACCTGACTCTGACAGGTGCAG |  |  |  |  |  |  |  |  |  |
|  |  | P: TGGAGAACACCCTCATCT |  |  |  |  |  |  |  |  |  |
|  | *prdx* | F: GGATCAACACCCCCAGGAA | 89.2 | 24.91 | 21.68 | 24.79 | 22.31 | 23.19 | 24.15 | 24.28 | - |
|  |  | R: GTCCTCCTTCAGCACTCCGTA |  |  |  |  |  |  |  |  |  |
|  |  | P: CCCTTGTGGCTGACCT |  |  |  |  |  |  |  |  |  |
|  | *il-1β* | F: TCAGGGTCTGGATCTGGAGG | 117.8 | 28.97 | 28.08 | - | 27.45 | - | - | - | 25.89 |
|  |  | R: CTCGGTTCCCATGGTAACCC |  |  |  |  |  |  |  |  |  |
|  |  | P: ACCCCATCACCATGC |  |  |  |  |  |  |  |  |  |
|  | *ifnγ* | F: TGTTTTCCCCAAGGACACGT | 116.3 | - | - | - | 30.01 | - | - | - | 31.01 |
|  |  | R: CCGATACACGTCCAGAACCA |  |  |  |  |  |  |  |  |  |
|  |  | P: TGAGCGGAGGGTGTT |  |  |  |  |  |  |  |  |  |
|  | *tnfα* | F: CTCAACTCTGTACGCACCGTG | 119.3 | 30.44 | 27.39 | - | 30.36 | - | - | - | - |
|  |  | R: TCTCCCTTCTCCAGGCTGAA |  |  |  |  |  |  |  |  |  |
|  |  | P: CGAGGCTGCGAGTGA |  |  |  |  |  |  |  |  |  |
|  | *mx* | F: GGAGGAGATTGAGGACCCCT | 105 | 28.83 | 24.78 | 28.89 | 25.81 | 29.22 | 29.27 | 27.31 | 28.51 |
|  |  | R: ATCACTGATACCCACCCCCA |  |  |  |  |  |  |  |  |  |
|  |  | P: TGAAGCCCAGGATGAA |  |  |  |  |  |  |  |  |  |
|  | *stat1* | F: CAGAAAGGCTTCCTGGAGGG | 100.9 | 27.87 | 24.35 | 28.42 | 25.54 | 26.50 | 26.94 | 23.71 | 24.53 |
|  |  | R: CTCTGTGGATGTGTGGGCAT |  |  |  |  |  |  |  |  |  |
|  |  | P: CGGGCCCTGTCACT |  |  |  |  |  |  |  |  |  |
|  | *ifn1* | F: CAGTATGCAGAGCGTGTGTCATT | 102.8 | 30.56 | 26.75 | 30.00 | 27.88 | 28.82 | - | 28.95 | 29.95 |
|  |  | R: TCTCCTCCCATCTGGTCCAG |  |  |  |  |  |  |  |  |  |
|  |  | P: CTGTGACTGGATCCGA |  |  |  |  |  |  |  |  |  |
|  | *il8* | F: TGGCCCTCCTGACCATTACT | 95 | - | 28.96 | - | 30.29 | 30.50 | 31.41 | 30.94 | 27.87 |
|  |  | R: GTCTCAATGCAGCGACATCG |  |  |  |  |  |  |  |  |  |
|  |  | P: ATGAGTCTGAGAGGCATG |  |  |  |  |  |  |  |  |  |
|  | *tapbp* | F: ACGGCAAGACTGACCGATTT | 99.2 | 29.14 | 30.95 | 29.34 | 28.91 | 29.76 | 30.72 | 30.07 | 30.48 |
|  |  | R: CACTTCAGCTTCCTGCAGGAT |  |  |  |  |  |  |  |  |  |
|  |  | P: AGCGGGGCTAGACTT |  |  |  |  |  |  |  |  |  |
| Endocrine disruption | *vtg1* | F: AAGTTCTGGGTAATGCTGGCC | 107.4 | - | 27.11 | - | - | 27.58 | - | - | 21.86 |
|  |  | R: CAAGACTGCATCAGCCTGGAC |  |  |  |  |  |  |  |  |  |
|  |  | P: TGCCCGTGTTTGGAA |  |  |  |  |  |  |  |  |  |
|  | *esr1* | F: GTGGAGGGTATGGCTGAGATCT | 103.2 | 29.69 | 25.14 | 29.10 | 29.29 | 31.23 | 31.57 | 34.88 | 31.88 |
|  |  | R: TCCTCAGGCTTCAGTTTAAGCAT |  |  |  |  |  |  |  |  |  |
|  |  | P: TGGCCACTGTGTCTC |  |  |  |  |  |  |  |  |  |
|  | *esr2a* | F: TTTGTGGACCTGTGCCTGTTC | 112.6 | 29.24 | 25.09 | 30.03 | 27.16 | 29.56 | 31.05 | 29.84 | 27.11 |
|  |  | R: CATGAGCCCTAGCATCAGCA |  |  |  |  |  |  |  |  |  |
|  |  | P: TTGGAGTGCTGCTGGT |  |  |  |  |  |  |  |  |  |
|  | *esr2b* | F: AACGAGGCCTGTCATTCCAG | 101.7 | 28.31 | 22.32 | - | 25.20 | 27.51 | 28.61 | - | 27.29 |
|  |  | R: TGTGTGAGAGCAGCATGAGGA |  |  |  |  |  |  |  |  |  |
|  |  | P: TCATCCCGCCTGGC |  |  |  |  |  |  |  |  |  |
|  | *ar* | F: AAGTGGTCAAGTGGGCCAAA | 90.7 | 28.85 | 25.04 | 26.88 | 24.70 | 27.04 | 27.53 | 26.88 | 26.83 |
|  |  | R: ACCCCCATCCATGAATGCT |  |  |  |  |  |  |  |  |  |
|  |  | P: TTGCCAGGTTTTCGGAAT |  |  |  |  |  |  |  |  |  |
|  | *cyp19a1b* | F: CGGACGACGTGAGACAGTGT | 83.7 | - | 28.80 | - | - | - | - | - | - |
|  |  | R: TCCTCATCTCCACCTCAGGG |  |  |  |  |  |  |  |  |  |
|  |  | P: TGATAGCAGCCCCAGACA |  |  |  |  |  |  |  |  |  |
| Circadian rhythm | *nr1d1* | F: TGTCCCGCGATGCTGTT | 102.6 | 27.83 | 25.95 | 27.20 | 22.87 | 26.56 | 27.21 | 29.68 | 27.18 |
|  |  | R: CCAGCATGCGCTGCTTCT |  |  |  |  |  |  |  |  |  |
|  |  | P: CTTTGGCAGAATACC |  |  |  |  |  |  |  |  |  |
|  | *nr1d2a* | F: TCGGATAGCGGCGATGA | 111.2 | 24.91 | 24.31 | 26.42 | 22.26 | 27.32 | 26.93 | - | 29.11 |
|  |  | R: GAATGTGCCCTGGTGACACA |  |  |  |  |  |  |  |  |  |
|  |  | P: AGGTTCCTCCTTCCAGG |  |  |  |  |  |  |  |  |  |
|  | *nr1d2b* | F: TCGGCATGTCCAAGGACTCT | 101 | 28.25 | 25.04 | 27.84 | 25.46 | 26.13 | 28.50 | 27.36 | 27.45 |
|  |  | R: CGCTGCTTCTCACGCTTTG |  |  |  |  |  |  |  |  |  |
|  |  | P: TGCGTTTCGGCCGCA |  |  |  |  |  |  |  |  |  |
|  | *arntl1a* | F: TCCAGCAGCCCGAGTAATG | 98.7 | 30.36 | 27.27 | 28.69 | 25.91 | 28.49 | 28.54 | 27.82 | 26.24 |
|  |  | R: GCCTCCAGAAGGCTCATGATC |  |  |  |  |  |  |  |  |  |
|  |  | P: CGAGGCAGCCATGG |  |  |  |  |  |  |  |  |  |
|  | *per1a/b* | F: CTGGACAACATTGCCTCAGAGTAC | 96.6 | 31.37 | 26.33 | 26.88 | 25.45 | 27.18 | 27.55 | - | - |
|  |  | R: GAAGGACACCGCCATGGA |  |  |  |  |  |  |  |  |  |
|  |  | P: CCCTCAAAAATACAGACACC |  |  |  |  |  |  |  |  |  |
|  | *per2* | F: GAGTACACCCTCAAAAACAATGACA | 112.4 | 31.19 | 27.64 | 29.30 | 29.93 | 30.80 | 31.11 | 30.98 | 28.70 |
|  |  | R: ATGTAGACGATCTTCCCCGTGAT |  |  |  |  |  |  |  |  |  |
|  |  | P: TGCGGTAGCGATATC |  |  |  |  |  |  |  |  |  |
|  | *cry1a/b* | F: CCGCCGGGACAAGGA | 98.3 | 27.15 | 25.52 | 30.76 | 22.64 | 26.39 | 29.44 | 27.37 | 27.12 |
|  |  | R: AGATGATTTCTACTCCGTGCTCTTC |  |  |  |  |  |  |  |  |  |
|  |  | P: TGGGCCGGTTAGC |  |  |  |  |  |  |  |  |  |
|  | *cry2* | F: TCCGCCTCTTCCTGAATGG | 142.5 | 28.85 | 25.53 | 27.76 | 25.28 | 28.51 | 27.70 | 25.00 | 27.31 |
|  |  | R: CCCACTACAGAGGCCTCAGTCT |  |  |  |  |  |  |  |  |  |
|  |  | P: TGGCATCTGTGCCC |  |  |  |  |  |  |  |  |  |
|  | *roraa/b* | F: TGCCGTGCCTTCGACTCT | 95.8 | 30.09 | 26.40 | 29.90 | 25.42 | 28.58 | 30.49 | 29.66 | 28.39 |
|  |  | R: CAGGCCCAGCATACTTTCCA |  |  |  |  |  |  |  |  |  |
|  |  | P: ACAACACAGTCTATTTCG |  |  |  |  |  |  |  |  |  |
|  | *clocka* | F: CGCAATGCAGCACCTGAA | 98.1 | 26.49 | 24.83 | 26.66 | 24.78 | 26.65 | 26.69 | 26.32 | 26.50 |
|  |  | R: ATGTTGGCCTCGATCATCCT |  |  |  |  |  |  |  |  |  |
|  |  | P: CTGGAGCAGAGGAC |  |  |  |  |  |  |  |  |  |
|  | *clockb* | F: CCTGGAGTCACTGGCCAAAT | 98.5 | 27.83 | 25.56 | 29.17 | 25.54 | 27.42 | 29.28 | 29.07 | 29.18 |
|  |  | R: CACGACTTCCCCTTCCCATA |  |  |  |  |  |  |  |  |  |
|  |  | P: CCACGAACACTTAATG |  |  |  |  |  |  |  |  |  |
|  | *npas2* | F: ACGCATGCTGCTCAACCA | 105.3 | 28.69 | 25.71 | 28.40 | 25.28 | 26.33 | 32.18 | - | - |
|  |  | R: CCTGGGAGCTAACACGGCTAT |  |  |  |  |  |  |  |  |  |
|  |  | P: CAGTGCAGACACTCGTA |  |  |  |  |  |  |  |  |  |
|  | *dbpa/b* | F: GAAGGGAGACCGTGCAAGTC | 102.4 | 26.53 | 27.09 | 27.79 | 25.71 | 34.22 | 30.74 | 35.03 | 33.12 |
|  |  | R: GGAGTCCTCATCCATGTCTGTGA |  |  |  |  |  |  |  |  |  |
|  |  | P: TGCGACGTTAAAGAC |  |  |  |  |  |  |  |  |  |
| Apoptosis | *casp3a/b* | F: AAAGGGATAGCTGCACAAGGG | 113.1 | 29.84 | 25.42 | - | 29.47 | 27.85 | 28.39 | - | - |
|  |  | R: TGTGTACATGTCAGCCGGAGG |  |  |  |  |  |  |  |  |  |
|  |  | P: CAGTCCCTCAGTGCAAA |  |  |  |  |  |  |  |  |  |
|  | *casp9* | F: GCCAGACAGTTGGTTCGAGAC | 86.5 | 31.97 | 28.13 | 30.54 | 28.32 | 28.19 | 31.40 | 33.29 | 31.82 |
|  |  | R: GGCTATGCTGCCCTTTCTCA |  |  |  |  |  |  |  |  |  |
|  |  | P: TCCCAGCTTTAATAGAG |  |  |  |  |  |  |  |  |  |
|  | *tp53* | F: CCTTTGAGGTGCGTGTGTGT | 116.1 | - | 26.83 | 27.75 | 25.66 | 26.96 | 28.53 | 27.86 | 26.36 |
|  |  | R: GGGTTGTCTCCTGCTGCTTCT |  |  |  |  |  |  |  |  |  |
|  |  | P: CTGGTCGAGACAGGAA |  |  |  |  |  |  |  |  |  |
|  | *pdcd10a/b* | F: AGGCTGAGAAGGAGAACCCAG | 90.2 | 28.31 | 23.61 | 29.37 | 24.70 | 25.74 | 28.89 | 28.39 | 28.91 |
|  |  | R: CACATCGTCTGCAGCCATTC |  |  |  |  |  |  |  |  |  |
|  |  | P: TGACCCAGGACATCAT |  |  |  |  |  |  |  |  |  |
|  | *chmp5a/b* | F: CCAAAGAGATGAAGGCGGC | 109.3 | 23.91 | 22.08 | 25.31 | 21.54 | 24.03 | 25.21 | 23.57 | 23.01 |
|  |  | R: TTGGCGTCCTCCATCATGT |  |  |  |  |  |  |  |  |  |
|  |  | P: AGGATCTCCAGGACCAG |  |  |  |  |  |  |  |  |  |
| Growth and metabolism | *ghr* | F: TAACCGGGAGCCACTTTGAC | 99.9 | 26.63 | 26.07 | 27.62 | 24.50 | 28.55 | 27.94 | 25.22 | 26.09 |
|  |  | R: ATTGACCTCACGGTACTGCACC |  |  |  |  |  |  |  |  |  |
|  |  | P: TGAGCTGGGAGCCG |  |  |  |  |  |  |  |  |  |
|  | *igf1* | F: TTCAAGAGTGCGATGTGCTGT | 97 | 31.09 | 23.14 | 29.60 | 25.62 | 25.73 | 27.27 | 33.61 | 29.29 |
|  |  | R: CGCCGAAGTCAGGGTTAGG |  |  |  |  |  |  |  |  |  |
|  |  | P: ACACCCTCTCACTGCT |  |  |  |  |  |  |  |  |  |
|  | *igf2* | F: ATGTGGAGGAGAACTGGTGGAC | 77.8 | 30.64 | 26.12 | 30.06 | 26.14 | 29.74 | 31.23 | 31.89 | 29.45 |
|  |  | R: CCTGCTGGTTGGCCTACTGA |  |  |  |  |  |  |  |  |  |
|  |  | P: TGCAGTTCGTCTGTGAAG |  |  |  |  |  |  |  |  |  |
|  | *igfbp1* | F: AACTGTGCGGAATCTACACGG | 98.5 | - | 29.61 | 30.97 | 25.81 | 28.26 | - | - | 29.31 |
|  |  | R: CTGGCTGCGAATAAGGGAGT |  |  |  |  |  |  |  |  |  |
|  |  | P: TGCACCCCAATACC |  |  |  |  |  |  |  |  |  |
|  | *igfbp2* | F: AGCTGCATGTCCTAAGCTTGC | 97.3 | 32.74 | 27.58 | - | 30.59 | 32.13 | - | - | 30.44 |
|  |  | R: GCCTTCTAACCGGGAACACA |  |  |  |  |  |  |  |  |  |
|  |  | P: TCAGGGAGCCCGGCT |  |  |  |  |  |  |  |  |  |
|  | *ampka1* | F: AAGTTTGAGTGCACCGAGGAG | 96 | 30.44 | 25.22 | 31.29 | 26.69 | 28.06 | 29.42 | 26.34 | 27.13 |
|  |  | R: GACATAATGCGGCGGTTGTC |  |  |  |  |  |  |  |  |  |
|  |  | P: CGCAACCACCACGAC |  |  |  |  |  |  |  |  |  |
|  | *ldhb* | F: CCCCCAACTGCACCCTTATT | 92.3 | - | 22.13 | 25.96 | 22.82 | 24.70 | 25.88 | 26.12 | 26.92 |
|  |  | R: AATCCGCTCAACTTCCACGT |  |  |  |  |  |  |  |  |  |
|  |  | P: CCAACCCAGTGGACGT |  |  |  |  |  |  |  |  |  |
|  | *pck1* | F: ATATGAGAACTGCTGGCTGGC | 90 | - | 30.55 | 28.54 | 28.97 | 31.38 | - | 31.63 | 29.28 |
|  |  | R: CACCTACTCGTGGAGACGGAA |  |  |  |  |  |  |  |  |  |
|  |  | P: CCCAGAGACGTGGCC |  |  |  |  |  |  |  |  |  |
|  | *fasn* | F: TGTGGGAGGTGTAGTCAAGCC | 106.5 | 27.50 | 21.40 | 27.56 | 25.56 | 24.86 | 25.63 | 28.26 | 26.10 |
|  |  | R: TCCCTGGGCCATGTATCTGA |  |  |  |  |  |  |  |  |  |
|  |  | P: AGGTGGAGGAGGCC |  |  |  |  |  |  |  |  |  |
|  | *cpt1a* | F: TACAGCTGGCCCAATTCAGG | 110.2 | 23.62 | 23.73 | 25.30 | 25.53 | 25.70 | 26.77 | 24.36 | 24.32 |
|  |  | R: AACCGTCTCTGTCCTACCCTCA |  |  |  |  |  |  |  |  |  |
|  |  | P: TCAATGACCCGGATGTT |  |  |  |  |  |  |  |  |  |
|  | *cpt1b* | F: GGACCAGTCCTGATGCCTTC | 100.4 | 24.17 | 23.60 | 25.50 | 21.96 | 23.90 | 25.64 | 24.57 | 24.87 |
|  |  | R: TCGAGGCCTCATACGTCAGAC |  |  |  |  |  |  |  |  |  |
|  |  | P: ACTGCAGCTGGCTCA |  |  |  |  |  |  |  |  |  |
|  | *cs* | F: TTGATTGCCAAGTTGCCGT | 112.1 | 23.00 | 21.19 | 21.92 | 20.61 | 21.22 | 21.75 | 23.23 | 21.60 |
|  |  | R: ATGTTGGCGAAGTTAGCGGA |  |  |  |  |  |  |  |  |  |
|  |  | P: CAGCAGCATTGGC |  |  |  |  |  |  |  |  |  |
|  | *ldha* | F: CGTCAAGTACAGCCCCAACG | 96.5 | 16.65 | 17.09 | 17.71 | 15.31 | 16.56 | 17.23 | 17.36 | 17.51 |
|  |  | R: GCCACGTAGGTCAGGATGTCA |  |  |  |  |  |  |  |  |  |
|  |  | P: TGCTGGTCGTCTCCAA |  |  |  |  |  |  |  |  |  |
|  | *lipea/b* | F: ACACTGGTCAAGGTGTTGCAGT | 111.5 | 29.56 | 27.63 | 29.31 | 26.70 | 29.22 | 27.88 | 29.71 | 27.95 |
|  |  | R: TGCAGTTAGCGGCAATGTAGC |  |  |  |  |  |  |  |  |  |
|  |  | P: CTTCTGCACATCATCCA |  |  |  |  |  |  |  |  |  |
|  | *lpl* | F: TGACAGCGCTGTACAAGAGGG | 105.8 | 27.19 | 22.90 | 26.15 | 24.67 | 27.01 | 26.31 | 30.23 | 25.59 |
|  |  | R: GGAGGTGAGGTAGTGCTGCTG |  |  |  |  |  |  |  |  |  |
|  |  | P: TGGACTGGCTGACACGG |  |  |  |  |  |  |  |  |  |
|  | *ctsd* | F: TTCACAGACATCGCCTGCTT | 109.4 | 23.04 | 21.94 | 24.05 | 21.41 | 23.26 | 23.79 | 21.55 | 23.85 |
|  |  | R: GGTACCCAGACAGACTGCCAG |  |  |  |  |  |  |  |  |  |
|  |  | P: CCACAAGTATAACGGTGCC |  |  |  |  |  |  |  |  |  |
| Detoxification | *cyp1a* | F: TACAGCTGGCCCAATTCAGG | 109.9 | 30.18 | 24.82 | 29.55 | 28.80 | 30.19 | 30.08 | - | 28.72 |
|  |  | R: AACCGTCTCTGTCCTACCCTCA |  |  |  |  |  |  |  |  |  |
|  |  | P: TCAATGACCCGGATGTT |  |  |  |  |  |  |  |  |  |
|  | *gstp1* | F: GGTGACAAGCCTTCGTTTGC | 91.2 | 26.96 | 21.40 | 25.02 | 22.92 | 23.18 | 24.21 | 24.69 | 25.31 |
|  |  | R: CAAAGCTCTTCAGGGAGGGG |  |  |  |  |  |  |  |  |  |
|  |  | P: TGAAGTGCTGCTCAAC |  |  |  |  |  |  |  |  |  |
|  | *gpx1a* | F: CCACCCCTTGTTTGTGTATCTCA | 79.3 | - | 25.16 | 28.50 | 25.66 | 26.79 | 28.47 | - | - |
|  |  | R: GGGCTCCACATGATGAACTTG |  |  |  |  |  |  |  |  |  |
|  |  | P: CCATTCCCCTCCGATG |  |  |  |  |  |  |  |  |  |
|  | *cat* | F: TGGGCCGCTACAACAGTACTG | 93.1 | 26.32 | 22.93 | 26.95 | 24.43 | 26.40 | 27.04 | 25.29 | 25.37 |
|  |  | R: CTCGTTCAGCACCTTAGTGAAGAA |  |  |  |  |  |  |  |  |  |
|  |  | P: CGTCACACAGGTGCGTA |  |  |  |  |  |  |  |  |  |
|  | *glul* | F: TGAAGTCATGCCTGCACAGTG | 95.1 | 28.09 | 24.17 | 31.29 | 26.18 | 28.78 | 26.18 | 27.86 | 27.21 |
|  |  | R: CACACCCGGTGGAGAATGA |  |  |  |  |  |  |  |  |  |
|  |  | P: CAGGTTGGCCCTTGT |  |  |  |  |  |  |  |  |  |
|  | *sod2* | F: GATGGCTGGGCTTTGACAA | 109.7 | 24.31 | 22.37 | 25.24 | 22.24 | 22.38 | 22.24 | 24.61 | 23.86 |
|  |  | R: CCTGCAGTGGGTCTTGATTAGG |  |  |  |  |  |  |  |  |  |
|  |  | P: AAGCTCCGTATCACAGC |  |  |  |  |  |  |  |  |  |
|  | *sod1* | F: CAACACCAACGGCTGTATGAGT | 97.9 | 33.44 | 21.01 | 24.15 | 23.56 | 22.93 | 23.82 | 23.72 | 22.76 |
|  |  | R: CCTCCGTGGGTCTTGTTGTG |  |  |  |  |  |  |  |  |  |
|  |  | P: CGGACCCCACTTCA |  |  |  |  |  |  |  |  |  |
|  | *nfe2l2a* | F: GGTGGCTACAGCGATTCAGAC | 108.2 | 27.57 | 24.58 | 28.18 | 25.49 | 27.15 | 28.84 | 25.85 | - |
|  |  | R: GAAAGAGGTGTCTGCAGGCC |  |  |  |  |  |  |  |  |  |
|  |  | P: AGTAACCCTGGGAGTGC |  |  |  |  |  |  |  |  |  |
| Hypoxia | *hif1a* | F: GCTGTGGGCTGAAGAGTGATC | 87.7 | 28.00 | 23.51 | 28.55 | 23.83 | 26.77 | 27.83 | 24.64 | 24.90 |
|  |  | R: CTGGGTTCTCCTTCAGCTGG |  |  |  |  |  |  |  |  |  |
|  |  | P: TCTCTGAGGCCTCCGAG |  |  |  |  |  |  |  |  |  |
|  | *epor* | F: TAAAGTGGCTCTGCTGCTCCA | 119.7 | 30.21 | 28.98 | 28.94 | 28.54 | 29.83 | 30.71 | - | - |
|  |  | R: CCAGAAGCAGGTGAGGTCTGA |  |  |  |  |  |  |  |  |  |
|  |  | P: CTGAGCCAGAGAATC |  |  |  |  |  |  |  |  |  |
|  | *vegfc* | F: AACCACACCGTGTGCAATTG | 93.9 | 33.27 | 27.40 | - | 26.63 | 27.26 | 29.73 | - | - |
|  |  | R: TGTGGTTCTTTGGGCAGGTT |  |  |  |  |  |  |  |  |  |
|  |  | P: ACTGGACGCCTACAGAC |  |  |  |  |  |  |  |  |  |
|  | *slc2a1a* | F: TGAGCATCGTGGCCATCTT | 123.6 | 24.66 | 24.44 | 27.21 | 25.26 | 27.58 | 25.60 | 24.82 | 24.45 |
|  |  | R: AACAGCTCAGCCACGATGAA |  |  |  |  |  |  |  |  |  |
|  |  | P: TTGGCCCGGGTCC |  |  |  |  |  |  |  |  |  |
|  | *mb* | F: ATGGTTCTGAAGTGCTGGGG | 117.3 | 25.20 | 25.14 | 30.46 | - | 30.97 | 29.83 | 28.41 | 29.10 |
|  |  | R: AAACAGACGGCTCAGAACCAG |  |  |  |  |  |  |  |  |  |
|  |  | P: AGGCTGACTACAACAAA |  |  |  |  |  |  |  |  |  |
|  | *hk1* | F: ATGCTGGAGGACATTCGCA | 126.6 | 28.90 | 27.15 | - | - | 30.48 | - | 27.35 | - |
|  |  | R: TTCGCCCATGTACATCCCA |  |  |  |  |  |  |  |  |  |
|  |  | P: AGGGGCTCTCTAAAC |  |  |  |  |  |  |  |  |  |
|  | *pfkm* | F: TGGTATCTACACCGGAGCCAA | 108.6 | 20.70 | 20.99 | 22.97 | 19.56 | 21.04 | 19.73 | 19.70 | 21.35 |
|  |  | R: TGCAGCATCATGGACACACTC |  |  |  |  |  |  |  |  |  |
|  |  | P: CCAGGGTCTGGTGGAT |  |  |  |  |  |  |  |  |  |
|  | *aldoaa* | F: CAACGGAGAGACCACCACTCA | 96.7 | 25.50 | 23.23 | 26.38 | 24.30 | 25.40 | 25.80 | 25.23 | 25.47 |
|  |  | R: ACGCCACTTAGCAAAGTCAGC |  |  |  |  |  |  |  |  |  |
|  |  | P: TGTACGAGCGGTGTGC |  |  |  |  |  |  |  |  |  |
|  | *eno1a* | F: TCCCTGCCTTCAACGTGATC | 93.7 | 21.77 | 19.47 | 21.90 | 19.37 | 20.21 | 21.27 | 23.16 | 22.80 |
|  |  | R: CCTCATGGCCTCCTTGAAGG |  |  |  |  |  |  |  |  |  |
|  |  | P: CGGAGGTTCCCACGCA |  |  |  |  |  |  |  |  |  |
|  | *pgk1* | F: AGATGATCATCGGTGGTGGC | 101.5 | 19.80 | 19.74 | 21.51 | 19.21 | 20.91 | 20.62 | 21.31 | 20.90 |
|  |  | R: CATACAGGGAGGTGCCGATC |  |  |  |  |  |  |  |  |  |
|  |  | P: CCTTCACCTTCCTCAATG |  |  |  |  |  |  |  |  |  |
| Endogenous control | *rpl7* | F: TCGTCATCAGGATCAGGGGT | 112.2 | 18.95 | 16.67 | 19.33 | 16.73 | 17.72 | 18.99 | 19.01 | 19.48 |
|  |  | R: GAAGATCTGACGCAGACGCA |  |  |  |  |  |  |  |  |  |
|  |  | P: CAAGGTGCGCAAGGT |  |  |  |  |  |  |  |  |  |
|  | *rps9* | F: TTCTCCCTGCGTTCACCATAC | 96.8 | 21.57 | 19.10 | 23.08 | 17.49 | 19.46 | 21.31 | 20.70 | 20.73 |
|  |  | R: GGCCCTTCTTGGCATTCTTT |  |  |  |  |  |  |  |  |  |
|  |  | P: CCCGGCCGTGTCAA |  |  |  |  |  |  |  |  |  |
|  | *ef1a* | F: GAAGCTTGAGGACAACCCCA | 106.7 | 19.18 | 18.68 | 19.61 | 17.01 | 18.95 | 18.94 | 19.24 | 19.96 |
|  |  | R: GAAGCTCTCCACACACATGGG |  |  |  |  |  |  |  |  |  |
|  |  | P: CCGCCATCATCGTCAT |  |  |  |  |  |  |  |  |  |
|  | *rpl13a* | F: CACTGGAGAGGCTGAAGGTGT | 104.1 | 21.33 | 19.19 | 21.16 | 18.50 | 19.91 | 20.83 | 20.83 | 20.96 |
|  |  | R: GTGGGCTTCAGACGGACAAT |  |  |  |  |  |  |  |  |  |
|  |  | P: CATGGTCGTACCTGCT |  |  |  |  |  |  |  |  |  |

Supplementary Table 4: Number and proportion (%) of the 112 qPCR assays that showed amplification using cDNA from gill, liver, and muscle tissue for the selected eight salmonid species.

|  | Gill | Percentage (%) | | Liver | Percentage (%) | Muscle | Percentage (%) |
| --- | --- | --- | --- | --- | --- | --- | --- |
| *S. salar* | 99 | 88 | 93 | | 83 | 93 | 83 |
| *S. trutta* | 100 | 89 | 94 | | 84 | 94 | 84 |
| *O. mykiss* | 92 | 82 | 90 | | 80 | 90 | 80 |
| *O. tshawytscha* | 103 | 92 | 96 | | 86 | 97 | 87 |
| *S. fontinalis* | 102 | 91 | 102 | | 91 | 102 | 91 |
| *S. alpinus* | 96 | 86 | 89 | | 79 | 90 | 80 |
| *C. clupeaformis* | 86 | 77 | 82 | | 73 | 81 | 72 |
| *C. hoyi* | 86 | 77 | 80 | | 71 | 88 | 79 |

Supplementary Table 5: The success rate of qPCR assays in 11 different biological pathways for cDNA from gill, liver and muscle tissues across all eight salmonid species.

| Biological function | Gill | | | Liver | | | Muscle | | |
| --- | --- | --- | --- | --- | --- | --- | --- | --- | --- |
|  | Total assays | Amplified assays | Success (%) | Total assays | Amplified assays | Success (%) | Total assays | Amplified assays | Success (%) |
| Stress response | 22 | 13 | 59 | 22 | 13 | 59 | 22 | 15 | 68 |
| Neural plasticity | 6 | 3 | 50 | 6 | 2 | 33 | 6 | 1 | 17 |
| Appetite regulation | 6 | 2 | 33 | 6 | 1 | 17 | 6 | 1 | 17 |
| Osmoregulation | 4 | 2 | 50 | 4 | 1 | 25 | 4 | 1 | 25 |
| Immune function | 12 | 9 | 75 | 12 | 7 | 58 | 12 | 5 | 42 |
| Endocrine disruption | 6 | 1 | 17 | 6 | 4 | 67 | 6 | 3 | 50 |
| Circadian rhythm | 13 | 9 | 69 | 13 | 9 | 69 | 13 | 10 | 77 |
| Apoptosis | 5 | 3 | 60 | 5 | 3 | 60 | 5 | 3 | 60 |
| Growth-metabolism | 16 | 12 | 75 | 16 | 10 | 63 | 16 | 12 | 75 |
| Detoxification | 8 | 6 | 75 | 8 | 6 | 75 | 8 | 6 | 75 |
| Hypoxia | 10 | 5 | 50 | 10 | 4 | 40 | 10 | 6 | 60 |
